# SARS-CoV-2 impacts the transcriptome and epigenome at the maternal-fetal interface in pregnancy

**DOI:** 10.1101/2022.05.31.494153

**Authors:** Lin Gao, Vrinda Mathur, Sabrina Ka Man Tam, Xuemeng Zhou, Ming Fung Cheung, Lu Yan Chan, Guadalupe Estrada-Gutiérrez, Bo Wah Leung, Sakita Moungmaithong, Chi Chiu Wang, Liona C. Poon, Danny Leung

## Abstract

During pregnancy, the maternal-fetal interface plays vital roles in fetal development. Its disruption is frequently found in pregnancy complications. Recent works show increased incidences of adverse pregnancy outcomes in COVID-19 patients; however, the mechanism remains unclear. Here, we analyzed the molecular impacts of SARS-CoV-2 infection on the maternal-fetal interface. Generating bulk and single-nucleus transcriptomic and epigenomic profiles from COVID-19 patients and control samples, we discovered aberrant immune activation and angiogenesis patterns in patients. Surprisingly, retrotransposons were dysregulated in specific cell types. Notably, reduced enhancer activities of LTR8B elements were functionally linked to the downregulation of Pregnancy-Specific Glycoprotein genes in syncytiotrophoblasts. Our findings revealed that SARS-CoV-2 infection induced significant changes to the epigenome and transcriptome at the maternal-fetal interface, which may be associated with pregnancy complications.

**One-Sentence Summary:** Pregnant COVID-19 patients show placental epigenetic and transcriptional changes, associated with adverse pregnancy outcomes.

## Main Text

The coronavirus disease 2019 (COVID-19) pandemic, caused by infection of the severe acute respiratory syndrome coronavirus 2 (SARS-CoV-2), has afflicted more than 520 million people with 6 million deaths globally (WHO website, accessed on May 24^th^, 2022). The situation worldwide remains dire. Evidence suggests that pregnant COVID-19 patients have worse clinical outcomes than similarly aged non-pregnant patients (*1, 2*). Indeed, pregnant patients show higher incidences of pneumonia, intensive care unit admission, need for invasive ventilation, need for extracorporeal membrane oxygenation, and death (*2–4*). Furthermore, SARS-CoV-2 infection during pregnancy is associated with adverse outcomes including preeclampsia, preterm birth, and stillbirth (*4–6*). Although recent reports have suggested that a subpopulation of trophoblasts in the placenta co-express SARS-CoV-2 receptor angiotensin-converting enzyme 2 (ACE2) and cofactor transmembrane serine protease 2 (TMPRSS2) in early pregnancy (*7, 8*), transplacental vertical transmission of the virus and placental infection are rarely detected (*2*). Therefore, it is likely that the COVID-19-related pregnancy complications result from maternal inflammatory response.

Pregnancy represents a unique immunological state, in which a dynamic balance between tolerating the fetus and defending against infections is maintained (*9*). This balance primarily occurs at the maternal-fetal interface, where the fetal placenta extensively invades and interacts with the maternal decidua (*10*). Interestingly, immune cells constitute about 40% of the maternal decidual cells during early pregnancy, with the majority being decidual natural killer (dNK) cells, decidual macrophages, and T cells (*11*). Unlike NK cells in the peripheral system, dNK cells have evolved specialized functions to promote endometrial decidualization. As demonstrated in both mice and humans, dNK cells interact with fetal HLA-expressing extravillous trophoblasts (EVT) to promote spiral arterial remodeling in the uterus, a critical process to ensure sufficient blood flow for the developing fetus (*12*). Decidual macrophages have an antigen-presenting role at the maternal-fetal interface. *In vitro* studies have found that they also produce vascular endothelial growth factor (VEGF) to promote angiogenesis (*11*), suggesting a role in placental angiogenesis. Immunosuppressive regulatory T (Treg) cells are also important for a successful pregnancy. A reduction in the numbers and functionality of Tregs at the maternal-fetal interface is associated with various placental pathology, including preeclampsia (*11*). Taken together, various fetal and maternal cells at the maternal-fetal interface work in concert to maintain the sophisticated immunological and angiogenic homeostasis during pregnancy. When this balance is broken, devastating consequences including placenta related adverse pregnancy outcomes will ensue.

Intriguingly, while viral infections frequently lead to adverse pregnancy outcomes, endogenous retroviruses (ERVs), which are ancient viral remnants within all metazoan genomes, possess critical roles in normal placental development (*13*). Retrotransposons, including ERVs, long interspersed nuclear elements (LINE) and short interspersed nuclear elements (SINE), are transposable elements that propagate in the genome by an RNA-mediated “copy-and-paste” mechanism and constitute more than 40% of the human genome (*14*). Although once considered as “junk DNA”, myriads of studies have shown that retrotransposons provide an abundant source of functional sequences for the host genome (*15*). Indeed, the most well-studied example of retrotransposon domestication has been found in the placenta, where the ERV-derived proteins, SYNCITINs, play an important role in the development of syncytiotrophoblasts (STs) (*16–18*). In addition, studies have shown that retrotransposons pervasively shape placental evolution, by supplying a diverse repertoire of *cis*-regulatory elements for endogenous genes (*19–23*). Notably, viral infections have been associated with aberrant derepression of retrotransposons, including SARS-CoV-2 (*24–26*). However, the effects of SARS-CoV-2 infection on these repetitive elements at the maternal-fetal interface remain unknown.

Currently, the underlying molecular mechanism of COVID-19-associated pregnancy complications is unclear, which necessitates further analysis. Given the complicated cell type composition at the maternal-fetal interface, single-nucleus-based, cell type-specific analysis is required. Moreover, although there have been several studies applying single-nucleus RNA-seq (snRNA-seq) on placental samples from COVID-19 patients to study the transcriptomic alterations (*27–29*), the epigenomic mechanism of the regulatory dysfunctions is still lacking. In this study, we investigated the cell type-specific molecular dysregulation at the maternal-fetal interface in SARS-CoV-2 infected pregnancies by mapping the transcriptomes and chromatin states at both bulk and single-nucleus resolutions with RNA-seq and ATAC-seq. We detected global transcriptomic and epigenomic alterations in patient samples, which highlighted immune activation and angiogenesis dysregulation. Strikingly, we discovered dysregulated retrotransposons, specifically LTR8B-derived enhancers, which were associated with the reduced expression of Pregnancy-Specific Glycoprotein (*PSG)* genes. Collectively, we generated extensive single-nucleus transcriptomic and epigenomic profiles of the maternal-fetal interface from COVID-19 and control patients, and our results highlighted the involvement of epigenetic regulation of *cis*-regulatory elements and retrotransposons in the maternal-fetal interface of COVID-19-related pregnancy complications.

## Results

### Multi-omic profiling of maternal-fetal interface in COVID-19 patients

To elucidate the molecular changes at the maternal-fetal interface upon SARS-CoV-2 infection during pregnancy, we assessed the transcriptome and chromatin states of patient and control samples at both bulk and single-cell levels (Fig. 1a). Maternal-fetal interface samples were collected from patients infected with SARS-CoV-2 during pregnancy, and from gestational age-matched control donors (<u>Table S1 and 2, fig. S1</u>). A total of 7 COVID-19 patients and 7 healthy pregnant donors were enrolled in this study. The demographic and clinical information for all subjects are listed in table S1-2 and fig. S1. COVID-19 patients had tested positive for SARS-CoV-2 by RT-qPCR (Ct <= 35) during late pregnancy (31.6-39.6 weeks). Among seven patients, six patients displayed mild symptom and one showed severe symptom with ICU admission and invasive ventilation (Cov6) (fig. S1). Three patients ended in pre-term delivery (gestational age < 37 weeks) and the remaining were full-term deliveries (Table S1). Maternal-fetal interface samples were collected immediately after delivery, and all the samples were defined as negative for SARS-CoV-2 by RT-qPCR (Table S3) and by N protein staining (<u>fig. S2a</u>).

**Fig. 1.**
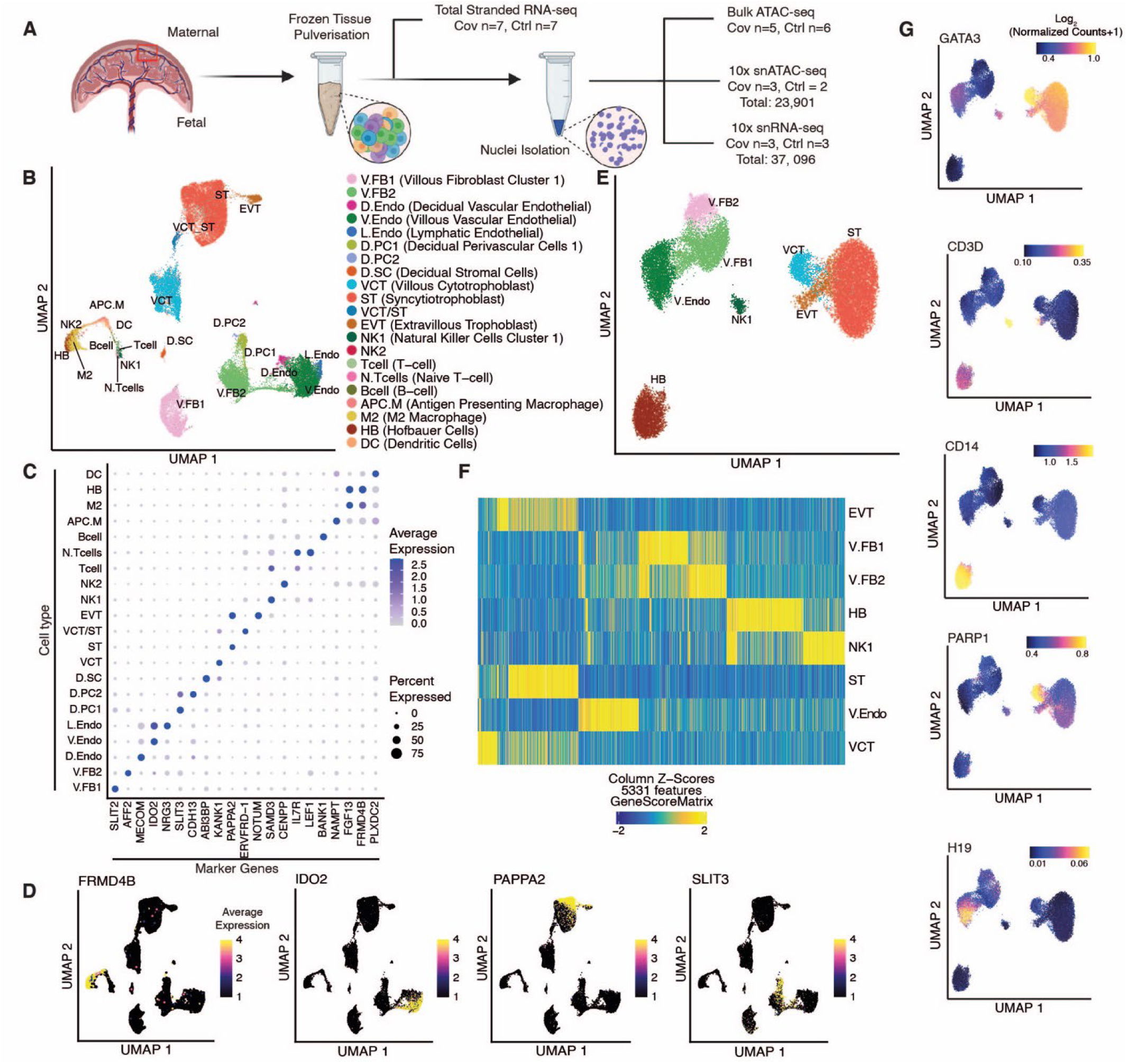
Multi-omic single-nucleus analysis of the maternal-fetal interface in COVID-19 patients. **(A)** A schematic of the experiment design. Frozen tissue samples are pulverized and subjected to RNA extraction for stranded total RNA-seq. The remaining tissues are processed for nuclei preparation and subsequent analysis by bulk ATAC-seq, snRNA-seq, and snATAC-seq. We then carry out integrative analysis on the multi-omic datasets derived from patient and control maternal-fetal interface samples. **(B)** UMAP of snRNA-seq showing 21 identified cell types. Each dot represents a single nucleus and its color corresponds to its annotated cell type. **(C)** Bubble plot of unique genes in each of the 21 cell types identified in snRNA-seq. The size of the bubbles represents the percentage of cells within a cluster that expresses the marker gene and the color of the bubble represents the average expression level of the marker gene calculated by Seurat. **(D)** Feature plots show the transcriptional levels of top marker genes and their corresponding cell type in snRNA-seq datasets. The colors of the dots represent the average expression level of the marker gene in each cell as calculated by Seurat. **(E)** UMAP of snATAC-seq with 8 cell types identified. Each dot represents a nucleus and the colors correspond to cell annotations transferred from the snRNA-seq. **(F)** Heatmap of the marker genes for all 8 cell types in snATAC-seq datasets. Each column represents the gene score of a marker gene and each row represents a cell type. The color of the heatmap indicates the z-score. **(G)** Feature plots show the gene score of top marker genes and their corresponding cell types in snATAC-seq datasets. The color of the dots represents the log_2_ normalized value.

We performed single-nucleus RNA-seq (snRNA-seq) on three patient samples and three control samples (10x Genomics) (<u>Table S4</u>). After quality control filtering, we analyzed a total of 37,096 nuclei (patients: n=18,782 and controls: n=18,314). Sequencing reads from each nucleus were mapped to both the human GRCh38/hg38 reference genome and the SARS-CoV-2 genome (severe acute respiratory syndrome coronavirus 2 isolate Wuhan-Hu-1, GenBank NC_045512.2 (*30*)). To delineate the cell type identities at the maternal-fetal interface, we performed unsupervised clustering with uniform manifold approximation and projection (UMAP), followed by cluster annotation with PlacentaCellEnrich (*31*), a tool for defining placental cell identities, and manual curation of the top expressed marker genes for each cell type (*32, 33*) (Fig. 1b, 1c, 1d, <u>fig. S2b</u>). We identified 10 major cell types at the maternal-fetal interface in patient and control samples, which aligned with the known cell categories at the maternal-fetal interface: trophoblasts (n=17,538), T cells (n=379), B cells (n=56), NK cells (n=288), macrophages (n=2,238), dendritic cells (n=326), endothelial cells (n=5,672), perivascular cells (n=962), fibroblasts (n=9,267), and stroma cells (n=368). We further subdivided these categories into 21 distinct cell types (Fig. 1b, <u>fig. S2c</u>). Concordant with our RT-qPCR and staining results, we detected no significant enrichment of viral transcripts in patient or control samples (<u>fig. S2d</u>). Consistent with the findings of a recent report (*27*), we noted low to undetectable expression of the *ACE2* and *TMPRSS2* genes in term samples from both patient and control samples (<u>fig. S2e</u>).

To investigate the cell type-specific alterations in gene regulation, we also mapped the chromatin accessibility at the maternal-fetal interface by single-nucleus ATAC-seq (snATAC-seq) (10x Genomics) (<u>Table S4</u>). In total, we generated chromatin state profiles for 23,901 individual nuclei from three patient and two control samples (patients: n=11,710 and controls: n=12,191) (Fig. 1e). We then performed label transfer from the snRNA-seq analyses to snATAC-seq clusters with ArchR (*34*). Based on cells’ chromatin state maps, we observed three major cell groups: trophoblasts, consisting of ST (n=9,024), EVT (n=994), and villous cytotrophoblasts (VCT) (n=1,467); immune cells, consisting of Hofbauer cells (HB) (n=2,935) and NK cells (n=344); and endothelial/fibroblasts cells, consisting of vascular endothelial cells (V.Endo) (n=3,373) and two fibroblast populations (V.FB1 and V.FB2) (n=1,585 and n=4,179, respectively) (Fig. 1e). Each cell type cluster showed distinct open chromatin patterns at transcriptional start sites (TSS) of marker genes and marker peaks defined by ArchR (Fig. 1f, <u>fig. S2f</u>). Moreover, we verified cell type annotations by investigating gene scores at known cell type-specific marker genes (Fig. 1g). Concordant with the snRNA-seq data, STs were the most abundant cell type among the profiled nuclei, likely due to their multinucleated property (<u>fig. S2c, 2g</u>).

To increase the depth of our analysis, we also conducted strand specific total RNA-seq and ATAC-seq on bulk patients and control samples (Fig. 1a <u>and</u> Table S4) (For RNA-seq, patients: n=7 and controls: n=7; for ATAC-seq, patients: n=5 and controls n=6). Comparing the reproducibility between the different data modalities, we surveyed the gene expression between aggregated snRNA-seq and bulk RNA-seq from the same biological samples and found a high degree of correlation between the two methods (<u>fig. S2h</u>). Similarly, the chromatin accessibility of active genes’ promoters assessed by aggregated snATAC-seq and bulk ATAC-seq datasets were consistent (<u>fig. S2i</u>). Taken together, we generated extensive single-nucleus and bulk multi-omic profiles of the maternal-fetal interfaces from control and SARS-CoV-2 infected pregnancies, which we utilized for subsequent integrative analyses.

### Extensive transcriptomic and epigenomic changes associated with SARS-CoV-2 infection

To investigate the cell type-specific molecular changes at the maternal-fetal interface upon SARS-CoV-2 infection, we identified the differentially expressed genes (DEGs) and differentially accessible regions (DARs) in patients from both bulk and single-nucleus datasets. We detected hundreds of DEGs from the bulk RNA-seq in patient samples (Fig. 2a) (upregulated: n=110, downregulated: n=456), suggesting extensive transcriptomic dysregulation upon SARS-CoV-2 infection. Notably, we discovered the downregulation of placenta development and pregnancy-related genes in COVID-19 patient samples, including the *PLAC1* gene and several members of *PSG* gene cluster, and the upregulation of interferon-related genes like *IFI6* and angiogenic factor gene *VEGFA* (Fig. 2a). Selected DEGs were validated by immunohistochemical staining (<u>fig. S3a</u>). From snRNA-seq data, we further defined cell type-specific DEGs for each cluster (Fig. 2b, <u>fig. S3b</u>). In both bulk and snRNA-seq analysis, we noted relatively high consistency of DEGs expression levels across individual patients (<u>fig. S3c</u>).

**Fig. 2.**
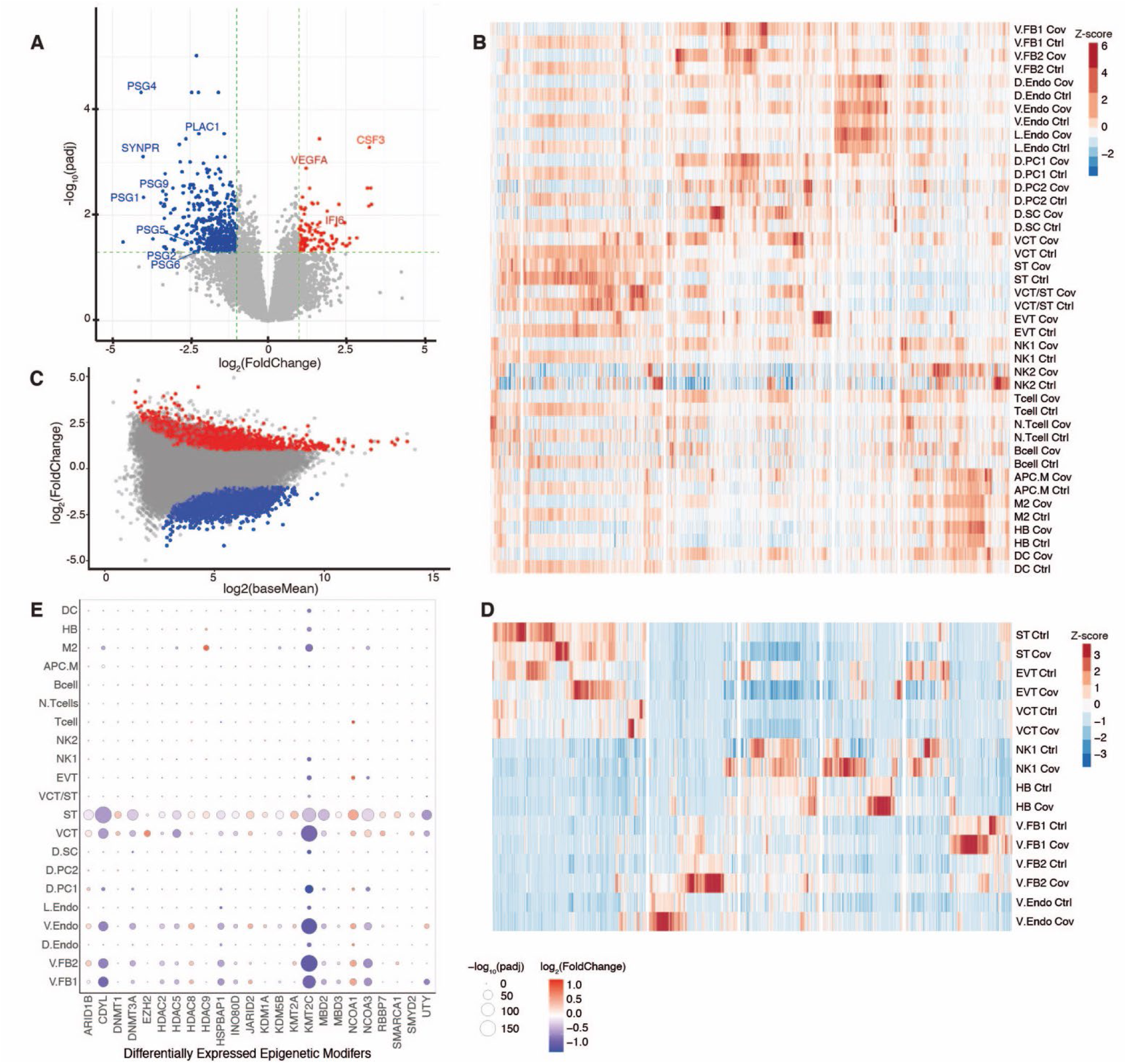
SARS-CoV-2 infection is associated with transcriptomic and epigenomic dysregulation. **(A)** Volcano plot shows significantly differentially expressed genes (DEGs) upon SARS-CoV-2 infection as measured by bulk RNA-seq. For each annotated gene, the negative log_10_-transformed two-tailed adjusted p-values (padj) from multiple testing using the Benjamini and Hochberg method are plotted against the log_2_-transformed fold change of fragments per kilobase per million reads (log_2_(FPKM) (patient/control). Upregulated (padj < 0.05 and log_2_(fold change) > 1; n=110) and downregulated (padj < 0.05 and log_2_(fold change) < −1; n=456) genes upon SARS-CoV-2 infection are labelled with red and blue, respectively. **(B)** Heatmap shows DEGs between patient and control samples in snRNA-seq datasets. Upregulated genes (padj <0.05 and log_2_(fold change) > 0.25) and downregulated genes (padj <0.05 and log_2_(fold change) < −0.25) are called by comparing patient and control cells within each cell type in the snRNA-seq analysis. **(C)** MA plot shows significantly differentially accessible regions (DARs) upon SARS-CoV-2 infection as measured by bulk ATAC-seq. For each peak, the negative log_2_-transformed fold change of reads per kilobase per million reads (log_2_(RPKM)) (patient/control) are plotted against the log_2_-transformed average signal. Increased (padj < 0.01 and log_2_(fold change) > 1; n=6,626) and decreased peaks (padj < 0.001 and log_2_(fold change) < −1; n=6,658) upon SARS-CoV-2 infection are labelled with blue and red, respectively. **(D)** Heatmap shows DARs between patient and control samples in snATAC-seq. Increased (padj<0.1 and log_2_(fold change) > 1) and decreased peaks (padj <0.1 and log_2_(fold change) < −1) are called by comparing patient and control cells within each cell type in the snATAC-seq datasets. **(E)** Bubble plot shows differentially expressed epigenetic modifiers between patient and control samples for each cell type by snRNA-seq. The size of the bubbles indicates negative log_10_ (padj) significance and the color indicates log_2_ (fold change) from the snRNA-seq differential analysis.

From the ATAC-seq profiles, we found thousands of DARs with gained or lost chromatin accessibility in patient samples (Fig. 2c, <u>fig. S3d</u>) (upregulated: n=6,626, downregulated: n=6,658), suggesting chromatin state alterations were also associated with SARS-CoV-2 infection. snATAC-seq also revealed cell type-specific DARs (Fig. 2d, <u>fig. S3e</u>). Motif analysis of regions with increased chromatin accessibility upon SARS-CoV-2 infection showed enrichment of CTCF motifs, suggesting these regions may be associated with novel *cis*-regulatory interactions (<u>fig. S3f</u>). Concordant with the downregulation of pregnancy-related genes in patient samples, DARs with reduced chromatin accessibility revealed the enrichment of placental transcription factor motifs, including the binding sites of TEAD2 and TEAD4 (<u>fig. S3f</u>).

Viral proteins are known to hijack the host’s epigenetic pathways for their propagation (*35*). It was recently reported that SARS-CoV-2 proteins could physically interact with epigenetic modifiers like histone deacetylases (HDACs) and bromodomain-containing proteins (*36*), making epigenetic modifiers potential therapeutic targets for COVID-19. Therefore, we analyzed the expression of epigenetic modifying enzymes within each cell type and identified 24 factors that were differentially expressed (Fig. 2e). We found genes encoding for HDAC enzymes to be dysregulated, including the downregulation of *HDAC2*, which was recently reported to be potentially inhibited by SARS-CoV-2’s non-structural protein 5 (NSP5) (*37*), suggesting that viral infection could also impact HDACs at the transcriptional level. In addition, DNA methyltransferase 1 (*DNMT1*) was upregulated in ST (Fig. 2e, <u>fig. S3g</u>), which could potentially cause DNA hypermethylation. This could be associated with the transcriptional downregulation and loss of chromatin accessibility observed in STs; however, additional experiments into the methylome are needed. Nevertheless, such transcriptomic and epigenomic alterations at the maternal-fetal interface may influence immunological and other responses in SARS-CoV-2 infected pregnancy. Therefore, we further investigated these processes in depth.

### Immune activation at the maternal-fetal interface upon SARS-CoV-2 infection

Cytokine storms and adverse immune-related outcomes have been reported in COVID-19 patients (*38*). We asked whether analogous immune dysregulation was present at the maternal-fetal interface in SARS-CoV-2 infected pregnancies. Consistent with a previous study that performed bulk RNA-seq in COVID-19 placenta samples (*27*), we also found the increased expression of interferon-related genes in COVID-19 patient samples. Concordantly, we found that upregulated genes were enriched with gene ontology (GO) terms relating to cytokine signalling and defense against viruses (<u>Fig. S4a</u>). Importantly, our snRNA-seq analysis revealed cluster-specific upregulation of interferon-related pathway genes in immune cells (Fig. 3a, 3b and 3c). Furthermore, regions with reduced chromatin accessibility upon SARS-CoV-2 infection were associated with GO terms related to negative regulation of immune suppression, in line with the immune activation in patient samples (<u>fig. S4d</u>). Indeed, analyzing snRNA-seq data, we found most of the detected interferon-inducible genes (IFI) and interferon-induced transmembrane genes (IFITM) to be significantly upregulated in at least one immune cell type (Fig. 3d). For example, the *IFI27* gene was upregulated in almost all immune cell types in patient samples, which was consistently observed in bulk RNA-seq datasets (Fig. 2a, 3d, 3e). It should be noted that the overexpression of *IFI27* transcripts in both the peripheral blood cells and plasma was reported to be a potential biomarker for SARS-CoV-2 infection (*39–41*). Our results suggest that *IFI27* upregulation could be a global immune cell response upon SARS-CoV-2 infection.

**Fig. 3.**
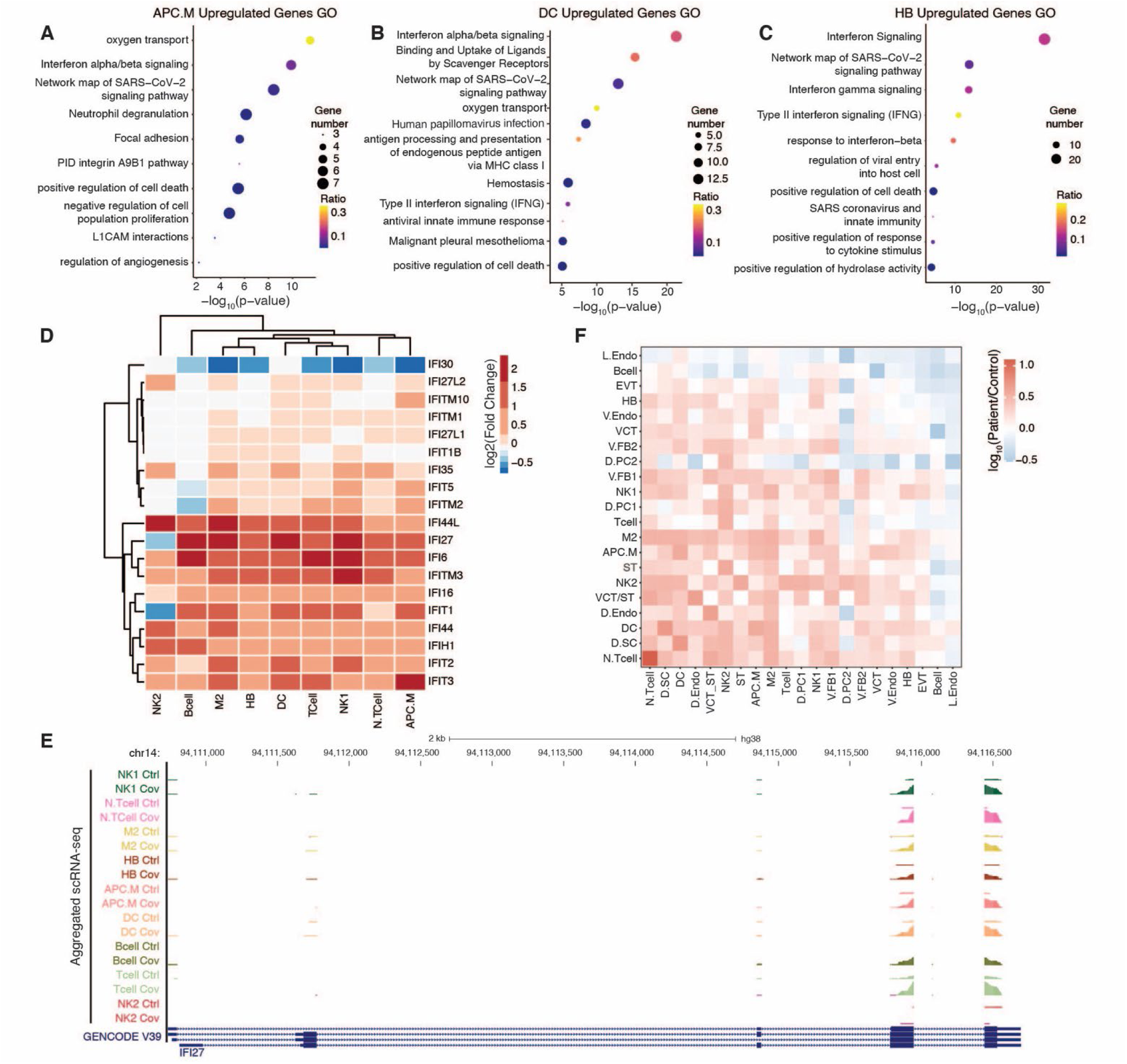
Immune activation at the maternal-fetal interface upon SARS-CoV-2 infection. **(A-C)** Bubble plots show Gene Ontology (GO) analysis of up-regulated genes from snRNA-seq in the **(A)** APC.M **(B)** DC, and **(C)** HB cell type clusters. The size and color of the bubble represent the number of upregulated genes and the ratio to total genes under each GO term, respectively. **(D)** Hierarchically clustered heatmap shows the log_2_ (fold change) of interferon gene expression in immune cell types as measured by snRNA-seq. **(E)** A genome browser screenshot of a genomic region surrounding the *IFI27* gene, which shows the patient and control pseudo-bulk tracks of snRNA-seq immune cell types are displayed as Reads Per Million mapped reads (RPM). The y-axis for all cell types ranges from 0 to 20. **(F)** Heatmap of CellPhoneDB receptor-ligand interactions, the color indicates the log_2_ (fold change) between the number of patient and control interactions calculated by CellPhoneDB.

To further characterize the dysregulation of immune-related pathways, we employed CellPhoneDB to investigate the alterations in cell type-specific receptor-ligand interactions within our snRNA-seq datasets (*42*). CellPhoneDB predicts the receptor-ligand interactions between cell populations by measuring the ligand expression in one cell cluster and the receptor expression in another cell cluster for each receptor-ligand pair (*42*). Interestingly, we detected significantly increased interactions within immune cell clusters and between immune clusters and other cell clusters, indicative of immune activation in SARS-CoV-2 infected pregnancies (Fig. 3f). Specifically, we found several chemokine production-related pairs to have increased interactions in patient samples (<u>fig. S4e, 4f)</u>. Taken together, we found a significant increase in immune response at the maternal-fetal interface characterized by the upregulation of interferon-induced genes and increased chemokine signalling upon SARS-CoV-2 infection.

### Dysregulation of angiogenesis pathways in COVID-19 pregnancies

Placental vasculature development and the remodelling of maternal spiral arteries in the maternal-fetal interface by invasive trophoblasts are vital for achieving a successful pregnancy (*43*). Abnormal angiogenesis is frequently observed in placental disorders and infections (*44–46*). Indeed, a recent study has reported increased levels of VEGF in placental villi by immunohistochemistry staining on COVID-19 samples, which is a feature of hypoxia and associated with preeclampsia (*47*). We asked whether SARS-CoV-2 infection would affect the angiogenesis pathway and what was the underlying molecular mechanism. Consistent with the previous report, we found that the upregulated genes in vascular endothelial cells and fibroblasts were related to blood vessel development and VEGF signalling (Fig. 4a, 4b). Concordantly, increased ATAC-seq peaks in these two cell types were associated with GO terms related to vasculogenesis and abnormal cell migration, suggesting epigenetic dysregulation of angiogenesis and blood vessel invasion pathways in patient samples (Fig. 4c, 4d). This finding was also observed in bulk ATAC-seq datasets (<u>fig. S4b</u>). Further, we analyzed the biological processes associated with COVID-19 sample-specific receptor-ligand interactions, which yielded GO terms including those related to angiogenesis processes (<u>fig. S4e</u>). For example, we detected several angiogenesis-related receptor-ligand pairs that were upregulated in COVID-19 patients, including VEGFA/FLT1 and FGF/FGFR (Fig. 4e). These receptor-ligand pairs are known to be involved in critical pathways for normal angiogenesis during placental development. Specifically, we found that gene *VEGFA* was upregulated in fibroblasts, which was accompanied by increased chromatin accessibility at its promoter region (Fig. 4f).

**Fig. 4.**
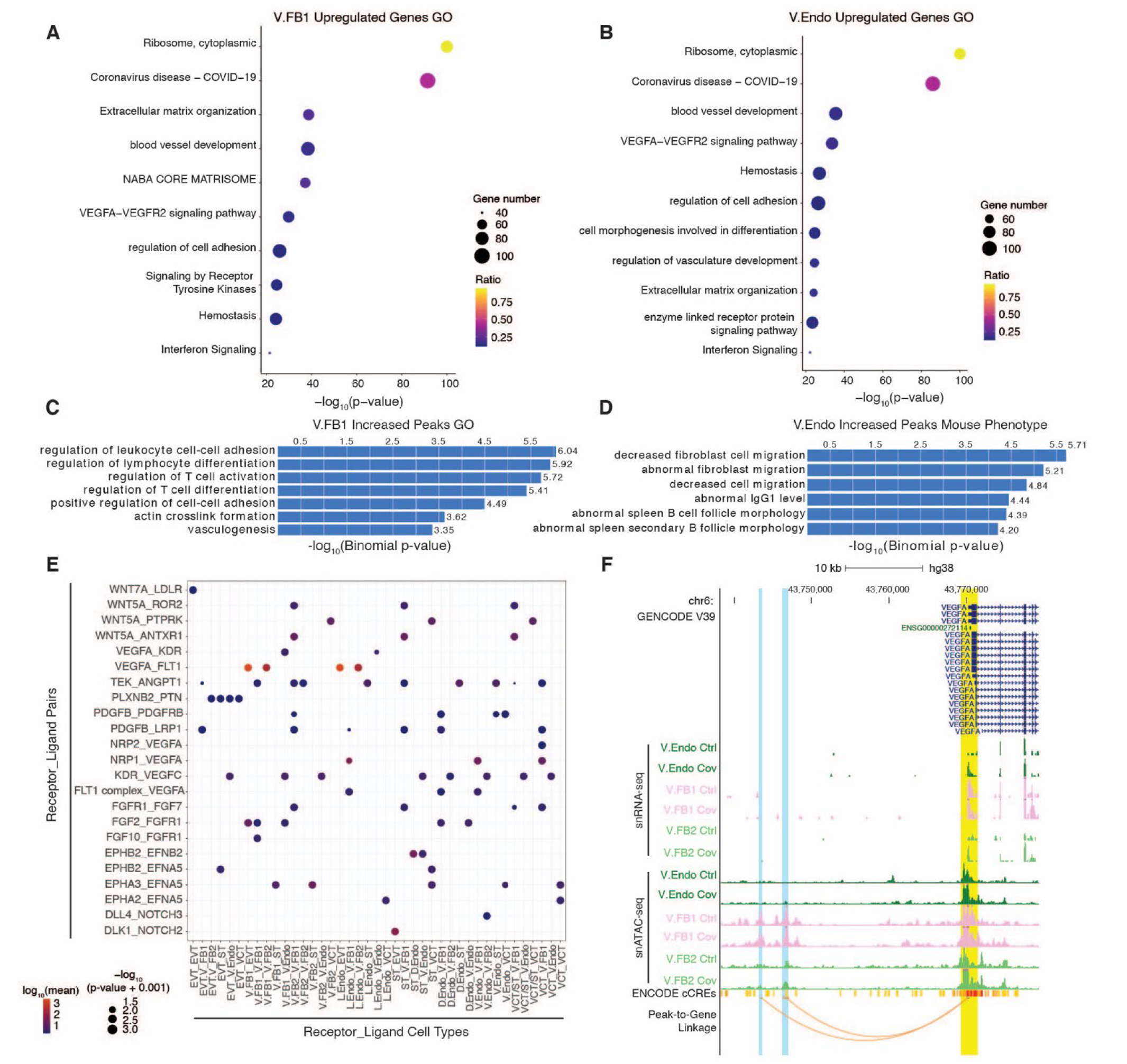
Dysregulation of angiogenesis pathways upon SARS-CoV-2 infection. **(A)** Bubble plot of top relevant GO terms for upregulated genes in V.FB1 cells from snRNA-seq. The size and color of the bubble represent the number of upregulated genes and the ratio to total genes under each GO term, respectively. **(B)** Bubble plot of top relevant GO terms for upregulated genes in V.Endo cells from snRNA-seq datasets. The size of the bubble indicates the number of genes found in our sample for each category and the color is the ratio of genes found to the total number of genes in the category. **(C)** Genomic Regions Enrichment of Annotation Tool (GREAT) analysis of increased peaks in V.FB1 cells from snATAC-seq showing top GO terms. The y-axis displays the negative log_10_ (binomial p-values). **(D)** GREAT analysis of increased peaks in V.Endo from snATAC-seq shows GO of single knockout mouse phenotype. The y-axis displays the negative log_10_ (binomial p-value). **(E)** Bubble plot of significant, patient specific receptor-ligand pairs under the GO term of angiogenesis. The color of the bubbles indicates the log_10_ mean, and the size of the bubbles shows the significance through negative log_10_ (p-value). **(F)** A genome browser screenshot of the *VEGFA* gene with patient and control tracks for V.Endo, V.FB1, and V.FB2 cell types from snRNA-seq and snATAC-seq displayed as pseudo-bulk RPM. The yellow shading highlights the promoter region of *VEGFA* and the blue shading highlights potential *cis*-regulatory enhancers called by peak-to-gene linkage (orange arcs). The y-axes for all cell types range from 0 to 2.

Next, we conducted peak-to-gene linkage analysis with ArchR, which integrated the chromatin accessibility states from snATAC-seq with gene expression from snRNA-seq to predict the potential enhancer-gene pairs (*34*). Intriguingly, we identified two candidate enhancers upstream of the *VEGFA* promoter, one of which coincides with a candidate *cis*-regulatory element (CRE) definition from the ENCODE consortium (Fig. 4f). Both candidate elements gained chromatin accessibility in fibroblasts (Fig. 4f), suggesting that transcriptional dysregulation of angiogenesis pathways may involve epigenetic alterations of associated *cis*-regulatory elements. Interestingly, we also discovered the upregulation of the *ENG* gene (<u>fig. S4g</u>), which encoded a glycoprotein endoglin, critical for blood vessel development. Notably, the elevated plasma concentration of soluble ENG (sENG) has been reported as a marker for preeclampsia, which is usually accompanied by increased circulating VEGF and Soluble fms-like tyrosine kinase-1 (sFLT1), suggesting pathological angiogenesis (*48, 49*). Concomitantly, we detected increased chromatin accessibility at its promoter and candidate enhancers in the corresponding cell types, which was bolstered by ENCODE candidate CRE definitions (<u>fig. S4g</u>). Collectively, our data suggested the dysregulation of angiogenesis in vascular endothelial cells and fibroblasts at the maternal-fetal interface in SARS-CoV-2 infected pregnancies, which involved differential activities of putative *cis*-regulatory elements.

### SARS-CoV-2 infection is associated with retrotransposon dysregulation in trophoblasts

Retrotransposons are repetitive sequences that constitute approximately 40% of the human genome. Studies have shown specific elements to be essential for mammalian placental development (*13, 50*). Moreover, viral infections have been suggested to associate with the derepression of retrotransposons in several diseases, including COVID-19 (*24–26*). However, the precise effects of SARS-CoV-2 infection on these repetitive elements, especially during pregnancy, remain unclear. Therefore, we analyzed the transcriptional and epigenomic states of these sequences in patient and control samples. Due to their repetitive natures, next-generation sequencing (NGS) reads from these elements will often be defined as multiple-aligned, which failed to be assigned to a specific locus. Hence, these reads are usually discarded, making it difficult to study retrotransposons with NGS-based assays. To circumvent this issue, we utilized our own iterative alignment approach termed Subfamily Assignment for Multiple Alignment (SAMA) (*51*), which rescues multiple-aligned reads and uniquely anchors them to retrotransposon subfamilies, enabling us to measure the transcriptomic and epigenomic changes with higher precision.

We first conducted differential analysis on bulk RNA-seq data from COVID-19 patient and control samples to identify retrotransposon subfamilies with an altered expression upon SARS-CoV-2 infection. Employing SAMA, we discovered three upregulated retrotransposon subfamilies and 21 downregulated subfamilies in patient samples, including the downregulation of the HERV17-int subfamily, from which the *SYNCITIN-1* gene was derived (Fig. 5a). Interestingly, most of the dysregulated subfamilies were ERVs (Fig. 5a). Many altered subfamilies detected from bulk data, such as MER65-int and HERVFH21-int elements, were similarly defined as dysregulated in snRNA-seq datasets, demonstrating the consistency across different platforms (<u>fig. S5a</u>). Strikingly, we also identified cell type-specific dysregulated subfamilies from single-nucleus analysis (Fig. 5b). For example, we found the LTR45B subfamily to be specifically downregulated in the ST cluster, suggesting a cell type specific regulatory mechanism for these elements (<u>fig. S5b)</u>. We then proceeded to investigate the expression of individual elements. We identified 926 upregulated elements and 6,884 downregulated elements in patient samples. As retrotransposons could provide CREs for host genes, we conducted GREAT analysis to test whether dysregulated elements were associated with particular groups of genes. Indeed, we found that upregulated elements were associated with GO terms for cell-cell adhesion genes, while downregulated elements were related to pregnancy genes (Fig. 5c, <u>fig. S5c</u>). Consistent with the subfamily analysis, we detected that individual HERV17-int, MER65-int and other elements were dysregulated (<u>fig. S5d</u>). We also identified individual dysregulated elements that were not defined in the subfamily analysis. For instance, we found several specific HERV3-int elements to be downregulated in patient samples, including an HERV3-int element derived gene (*ERV3-1)* (Fig. 5d, <u>fig. S5d</u>). Intriguingly, further analysis with snRNA-seq showed the downregulation of this element was restricted to several cell types, primarily in STs and VCTs (<u>Fig. 5d</u>). Notably, ERV3 class elements are known to be highly expressed in the placenta and their decreased expression has been reported to be associated with pregnancy disorder such as intrauterine growth restriction (*52*). Our finding suggests pregnancy-related ERVs are dysregulated in the maternal-fetal interface in SARS-CoV-2 infected patients.

**Fig. 5.**
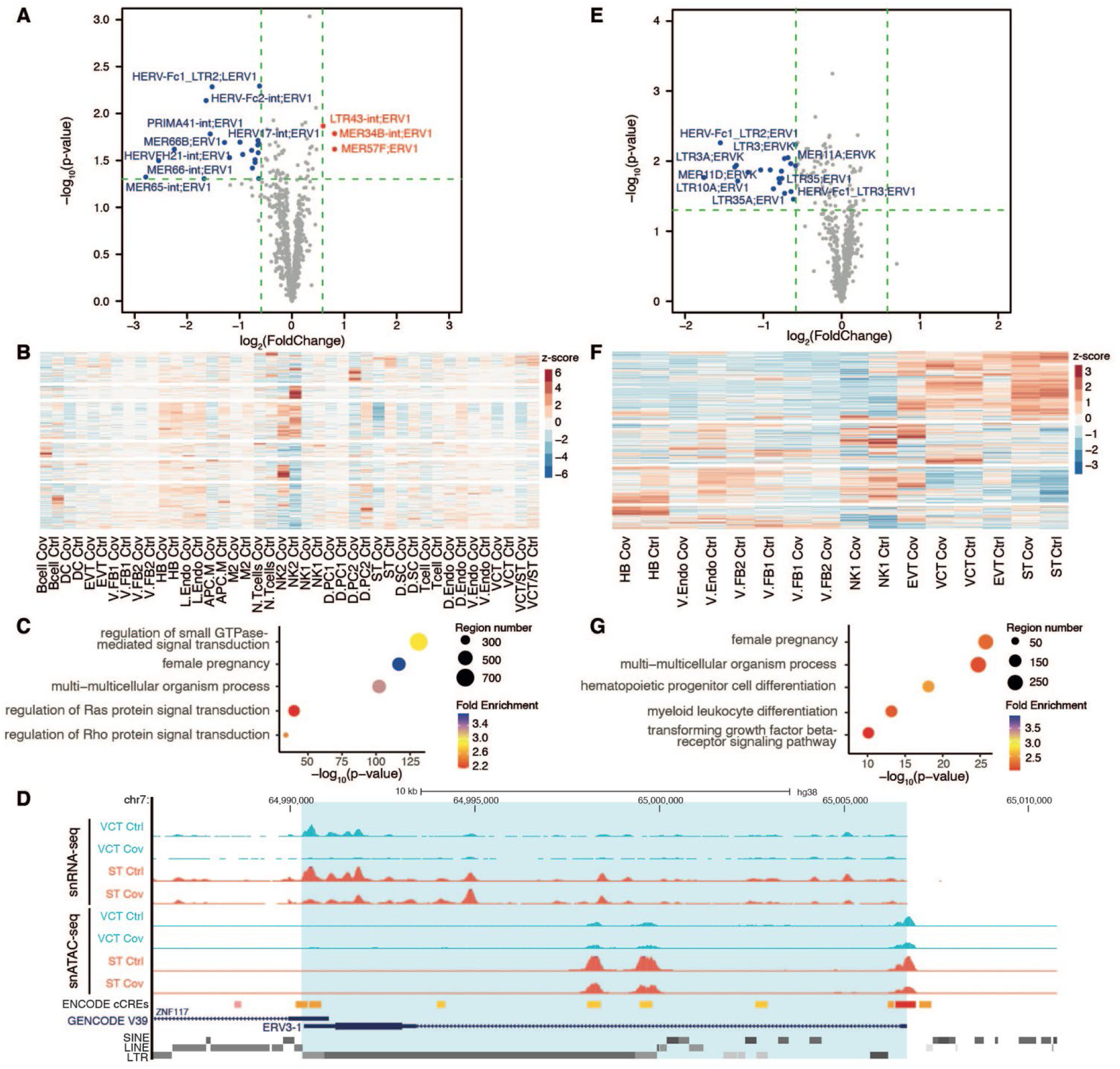
Dysregulation of retrotransposons in SARS-CoV-2 infected pregnancies. **(A)** Volcano plot showing differentially expressed retrotransposons subfamilies between patient and control samples as measured by bulk RNA-seq. For each subfamily, the negative log_10_ (padj) is plotted against the log_2_ (fold change) of RPKM (patient/control). Significantly upregulated subfamilies (padj < 0.05 and fold change > 1.5; n = 3) and downregulated subfamilies (padj < 0.05 and fold change < 0.66; n = 21) are labelled with red and blue, respectively. **(B)** Heatmap showing expression of retrotransposon subfamilies in distinct cell types as determined by snRNA-seq. Color scale represents RPM-calculated z-score. **(C)** GREAT analysis of significantly downregulated individual retrotransposons from bulk RNA-seq analysis of patient samples shows enrichment of GO terms including female pregnancy. **(D)** A genome browser screenshot shows an example of upregulated retrotransposon ERV3-1 in ST and VCT cells. snRNA-seq and snATAC-seq datasets are displayed as pseudo-bulk RPM, with y-axis ranging from 0 to 10 and 0 to 5, respectively. **(E)** Volcano plot showing retrotransposons subfamilies with differential chromatin accessibility as measured by bulk ATAC-seq. For each subfamily, the negative log_10_ (padj) are plotted against the log_2_ (fold change) of FPKM (patient/control). Retrotransposon subfamilies with significantly increased (padj < 0.05 and fold change > 1.5; n = 0) and decreased (padj <0.05 and fold change < 0.66; n = 19) signals are labelled with red and blue, respectively. **(F)** Heatmap showing chromatin accessibility of retrotransposon subfamilies across cell types as determined by snATAC-seq. Color scale represents RPKM-calculated z-score. **(G)** GREAT analysis of individual retrotransposons with decreased chromatin accessibility from bulk ATAC-seq shows enrichment of GO terms including female pregnancy.

We then investigated whether retrotransposon-derived CREs were also dysregulated in patient samples by analyzing their chromatin accessibility. Using SAMA, we identified retrotransposon subfamilies with altered bulk ATAC-seq signals in patient samples. Surprisingly, all differentially regulated retrotransposon subfamilies showed decreased chromatin accessibility, and no subfamilies gained chromatin accessibility (Fig. 5e). Furthermore, from snATAC-seq datasets, we observed cell type-specific dysregulation of retrotransposons, which were obfuscated in the bulk data (Fig. 5f). For example, although we did not detect subfamilies with increased accessibility from bulk ATAC-seq, we found several subfamilies including the HERV-K LTR5, which significantly gained chromatin accessibility in STs (<u>fig. S5e</u>). Moreover, while HERV17-int subfamily had decreased chromatin accessibility in most clusters, the change in ATAC-seq signals and the degree of significance were most substantial in trophoblasts clusters (<u>fig. S5f</u>). We then overlapped ATAC-seq peaks with individual element annotations from the RepeatMasker (GRCh38/hg38). Considering individual retrotransposons, we discovered elements with altered chromatin accessibility that were not present in subfamily analysis (<u>fig. S5g</u>). Elements with increased chromatin accessibility showed enrichment of transcription factor motifs relating to the regulation of the NF-*κ*B signalling pathway (<u>fig. S5h</u>), suggesting their involvement in the activation of innate immunity in COVID-19 pregnancies. Interestingly, we again found GO terms relating to female pregnancy that was associated with retrotransposons with decreased chromatin accessibility (Fig. 5g). Elements with decreased chromatin accessibility in the ST cells cluster were enriched with binding motifs for TFs with known placental functions, including GRHL2, TEAD4 and JUNB (<u>fig. S5i</u>). These results suggest that changes in retrotransposon activities at the maternal-fetal interface upon SARS-CoV-2 infection may be involved in the dysregulation of important pregnancy genes.

### Expression of pregnancy genes is reduced concomitantly with loss of chromatin accessibility at intronic retrotransposons

As mentioned, we observed the downregulation of pregnancy-related genes including the *PSG* at the maternal-fetal interface in SARS-CoV-2 infected pregnancies (Fig. 2a). The human genome contains ten *bona fide PSG* genes, *PSG1*-*9* and *PSG11*, which are clustered on chromosome 19q13. *PSG10* is classified as a pseudogene (*53*). *PSG* genes are expressed at high levels during pregnancy and are involved in the immune-modulation and angiogenesis processes (*53*). Notably, reduced serum PSG level is associated with pregnancy complications, including intrauterine growth restriction, pregnancy loss, and preeclampsia (*54–56*). However, the precise regulatory mechanism of the *PSG* gene cluster remains poorly understood. We found that 8 out of 10 *PSG* genes were downregulated in the ST cluster in patient samples (Fig. 6a, b, fig. <u>S6a</u>), which were similarly detected in bulk RNA-seq data (<u>fig. S6b</u>). We further validated the changes in transcriptional and protein levels by RT-qPCR and immunohistochemical staining, respectively (Fig. 6c, 6d). Intriguingly, we observed that the downregulation of *PSG* genes occurred concomitantly with decreased chromatin accessibility of intronic LTR8B elements within the said genes (Fig. 6e, <u>fig. S6c</u>). Each of the downregulated genes harboured an LTR8B element with consistent loss of ATAC-seq signals. Notably, these intronic LTR8B elements showed trophoblast-specific open chromatin states (<u>fig. S6d</u>). We postulate that these elements may be putative trophoblast-specific regulatory elements for the *PSG* genes.

**Fig. 6.**
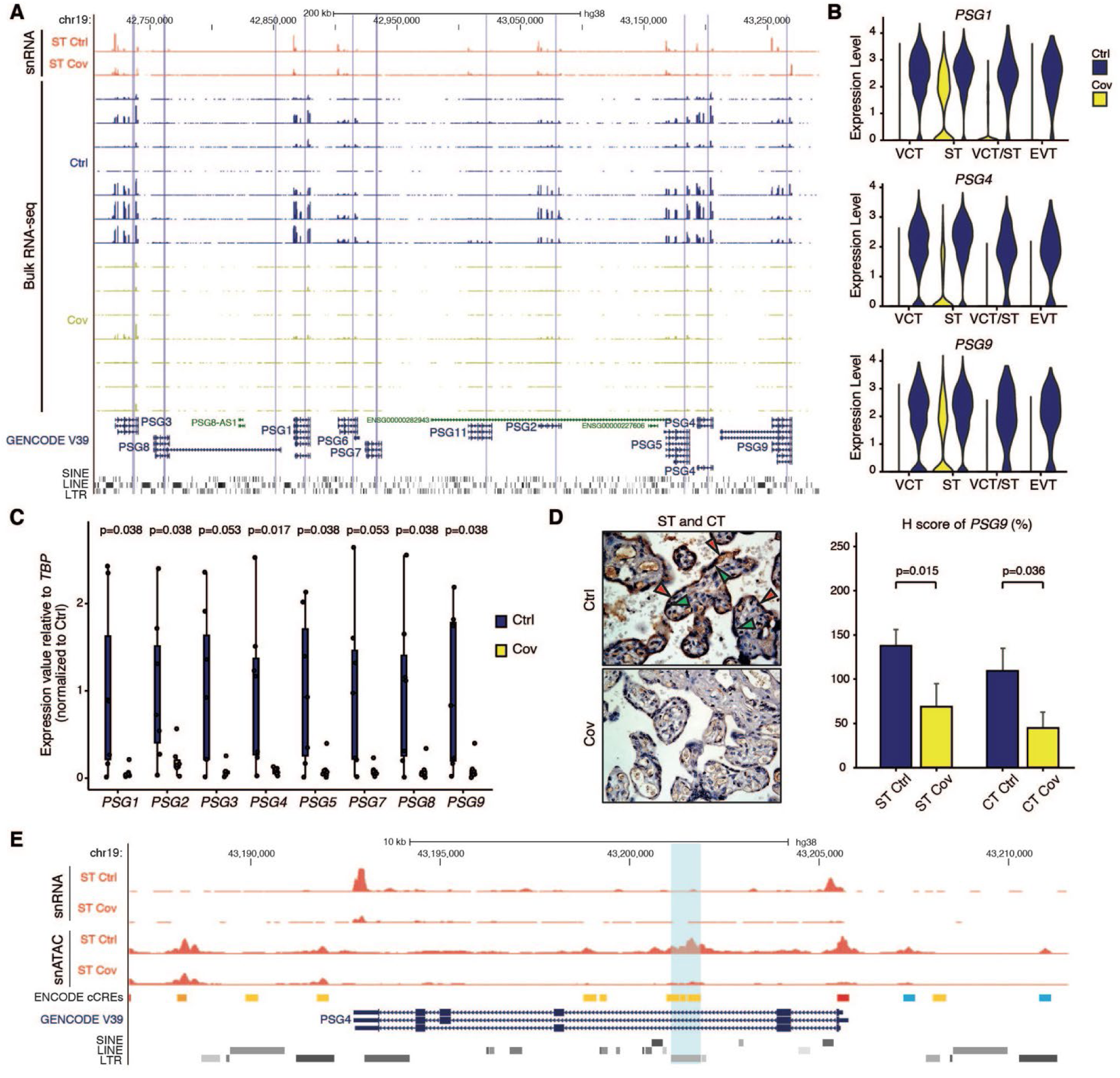
Reduction of pregnancy gene expression was associated with loss of chromatin accessibility at intronic LTR elements. **(A)** A genome browser screenshot of *PSG* gene cluster shows decreased expression *en masse* in both the single nucleus and bulk RNA-seq datasets in patient samples. Purple shading denotes the location of LTR8B elements within each *PSG* gene. Pseudo-bulk snRNA-seq and bulk RNA-seq datasets are displayed as RPM, with y-axis ranging from 0 to 350 and 0 to 100, respectively. **(B)** Violin plots showing decreased expression of *PSG1* (top), *PSG4* (middle), and *PSG9* genes (bottom) in different trophoblast cells from snRNA-seq in patient versus control samples. Expression values are displayed as normalized counts. **(C)** RT-qPCR of 8 of the *PSG* genes show decreased expression in patient samples. Results are displayed as expression values relative to control samples. Each dot represents one sample. P-values represent significance as calculated by the student T-test. **(D)** Immunohistochemistry staining (left) and H score (right) of PSG9 protein shows decreased protein levels in patient samples. Red arrows indicate ST cells and green arrows indicate CT cells. P-value is calculated by the student T-test. **(E)** A genome browser screenshot showing decreased expression of the *PSG4* gene and increased chromatin accessibility at intronic LTR8B element in patient samples. Pseudo-bulk snRNA-seq and snATAC-seq datasets are displayed as RPM, with y-axis ranging from 0 to 100 and 0 to 5, respectively. The blue shade highlights LTR8B elements within the *PSG4* gene.

### LTR8B elements function as enhancers in placental cells

Given their chromatin states, we sought to investigate the enhancer potentials of the intronic LTR8B elements in regulating the *PSG* genes. We analyzed publicly available RNA-seq and ChIP-seq datasets generated from specific primary placenta cell types, found in the International Human Epigenome Consortium (IHEC) data repository (*57, 58*). We found that the high expression of *PSG* genes in ST cells was accompanied by enrichment of active enhancer histone modifications (H3K27ac and H3K4me1) and low levels of active promoter signatures (H3K4me3). As expected, repressive modifications, such as H3K9me3 and DNA methylation, were depleted at these retrotransposons, supporting the model that these LTR8Bs function as enhancers to regulate *PSG* genes’ expression (Fig. 7a). Notably, the activation of these intronic LTR8Bs was unlikely to be simply a positional effect. For instance, while the LTR8B element located in the intronic region of *PSG8* was enriched with active histone marks, another ERV1 cluster in the same intron of *PSG8* was marked by H3K9me3 (Fig. 7a), suggesting its activity was related to its sequence features. Therefore, we confirmed the enhancer functions of LTR8B elements by luciferase assay, in which most elements demonstrated strong enhancer activities (Fig. 7b, <u>fig. S7a</u>).

**Fig. 7.**
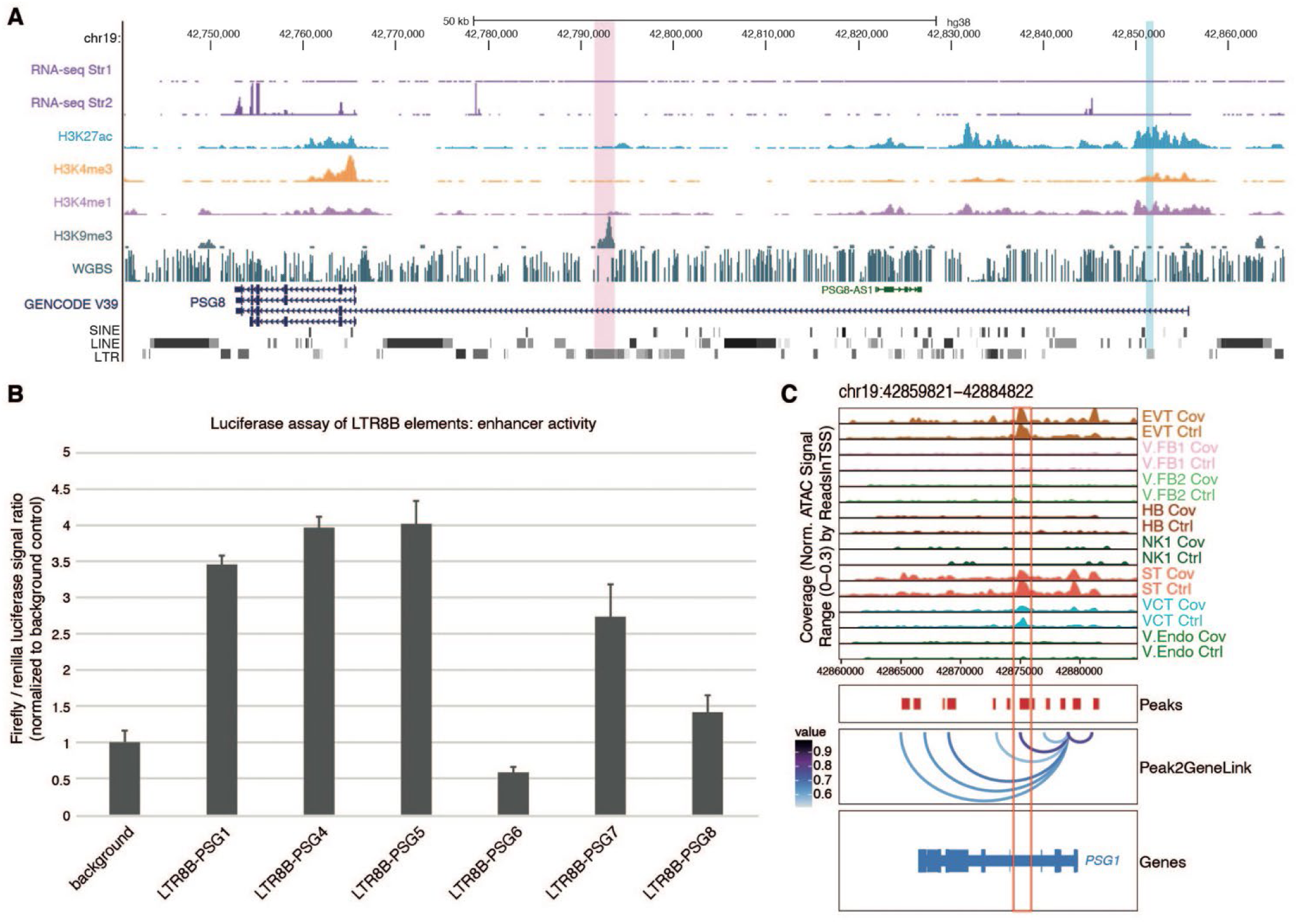
LTR8B elements function as enhancers in trophoblast cells. **(A)** A genome browser screenshot showing the transcriptional and epigenetic states of the *PSG8* gene in primary STs. Datasets are obtained from the IHEC data repository (*57, 58*). All datasets are displayed as RPM, with y-axis ranging from 0 to 200 for RNA-seq, 0 to 100 for H3K27ac ChIP-seq, 0 to 250 for H3K4me3 ChIP-seq, 0 to 150 for H3K4me1, 0 to 20 for H3K9me3 ChIP-seq, and 0 to 1 for Whole Genome Bisulphite sequencing (WGBS). The pink shading highlights the intronic ERV1 element and the blue shade highlights LTR8B within *PSG8*. **(B)** Luciferase assay of individual LTR8B elements within different *PSG* genes suggests that the majority possesses enhancer activity. Y-axis value represents Firefly/renilla luciferase signal ratio normalized to background control. **(C)** Peak-to-gene linkage analysis indicates a potential interaction between the *PSG1* promoter and its intronic LTR8B (red box). Datasets are displayed as pseudo-bulk signals of each cell type from snATAC-seq.

We then explored the functional relationship between the LTR8B-derived enhancers and *PSG* genes’ expression. We found the degree of chromatin accessibility at intronic LTR8B elements was significantly correlated with the expression levels of corresponding PSG genes (<u>fig. S7b</u>), supporting the enhancer function of these elements. Moreover, peak-to-gene linkage analysis from snATAC-seq data identified significant linkages between *PSG* promoters and their intronic LTR8B elements (Fig. 7c, <u>fig. S7c</u>). To elucidate the precise targets of LTR8B enhancers, we employed the human expanded potential stem cells (EPSC) derived trophoblast stem cells (TSCs) as a model system (*59*). We found that these cells had high expression of *PSG*s and concordant enrichment of active enhancer histone marks at the intronic LTR8B elements (<u>fig. S7d</u>). To determine the higher-order chromatin interactions of these loci, we performed Hi-C in TSCs. Interestingly, our data revealed strong interactions between the intronic LTR8B elements and multiple *PSG* gene promoters (Fig. 8a), suggesting that these retrotransposons formed a regulatory network for the *PSG* gene cluster.

**Fig. 8.**
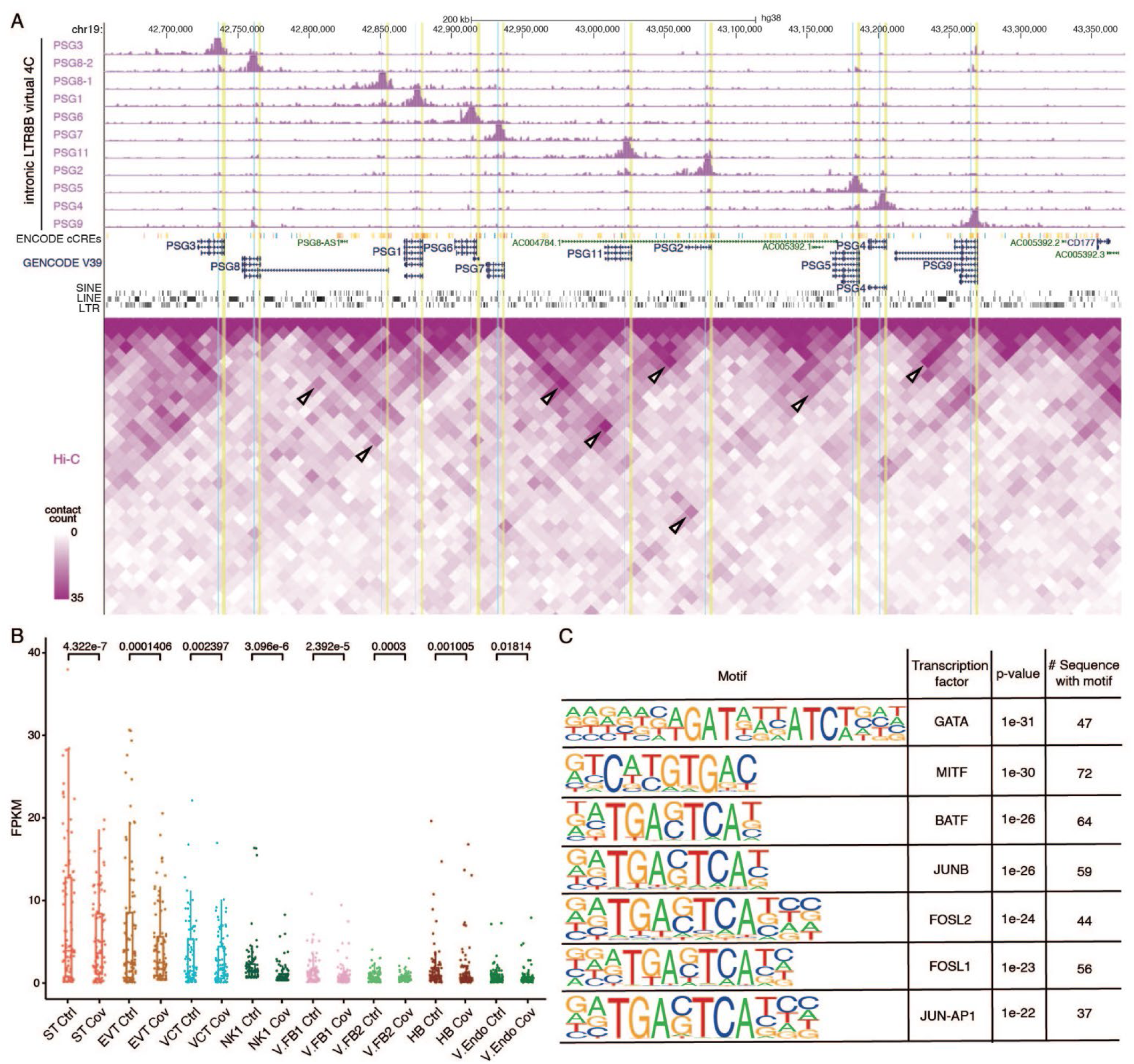
Intronic LTR8B elements interact with *PSG* gene promoters. **(A)** A screenshot of the virtual 4C results (top) and Hi-C interaction frequency from trophoblast stem cells (TSCs) (bottom) at the *PSG* cluster shows the interaction of *PSG* gene promoters and intronic LTR8B elements. Virtual 4C shows interaction frequencies using each intronic LTR8B element as an anchor and datasets are displayed as RPKM, with y-axis ranging from 0 to 20000. The intronic LTR8B elements and the promoters of *PSGs* are highlighted with blue and yellow shading, respectively. White arrowheads in the Hi-C heatmap point to interactions within the *PSG* cluster, indicating a high interaction frequency between intronic LTR8B and *PSGs* promoters. **(B)** Boxplot showing the pseudo-bulk FPKM signal of all active LTR8B elements defined by co-accessibility analysis (n=91) in each cell types from snATAC-seq. Results suggested that active LTR8B elements have higher snATAC-seq signals in trophoblast cells (ST, EVT, and VCT) and exhibited significantly decreased chromatin accessibility upon SARS-CoV-2 infection. One-tailed paired T-test was applied to measure the significance. Each dot represents one active LTR8B element. **(C)** HOMER motif analysis of active LTR8B elements defined by co-accessibility analysis (n=91) reveals significant enrichment of transcriptional factor binding motifs, including those of GATA, FOSL, and JUN-AP1 families of TFs.

To determine whether LTR8B elements broadly functioned as enhancers in the placental transcriptional regulation and whether they were similarly affected by SARS-CoV-2 infection, we interrogated other elements within the subfamily. LTR8B subfamily contains a total of 1,600 individual elements. Chromatin accessibility of the LTR8B subfamily was particularly high in STs compared to other cell clusters. Strikingly, the accessibility of the whole subfamily was significantly decreased in patient ST cells (<u>fig. S7e</u>). It should be noted that other LTR8B elements outside of the *PSG* gene cluster also possessed open chromatin states in STs (<u>fig. S7f</u>). We further defined 91 LTR8B elements as putative genic enhancers from peak-to-gene linkage analysis. We found that active LTR8Bs showed relatively high chromatin accessibility in trophoblast cell clusters, including STs, EVTs, and VCTs (Fig. 8b). Concomitant with SARS-CoV-2 infection, a marked decrease in ATAC-seq signal was detected in all the cell types, with the most significant change in STs (Fig. 8b). Moreover, these active elements were enriched with motifs for TFs related to placental development, including GATA, FOSL1, FOSL2, and AP-1, suggesting that they function as important *cis*-regulatory elements during placenta development (Fig. 8c). For example, an LTR8B element with active enhancer properties was predicted to be a potential enhancer for gene *STS*, which was downregulated in STs (<u>fig. S7g</u>). This gene encodes for protein steroid sulfatase, an enzyme in the estrogen synthesis pathway and is highly expressed in ST cells (*60*). Placental STS deficiency is known to lead to low estrogen levels, prolonged pregnancy, and dysfunctional labour (*61*). Consistent with the downregulation of *STS*, the LTR8B element showed lower ATAC-seq signals in COVID-19 samples (<u>fig. S7g</u>). This LTR8B element was enriched with active enhancer marks H3K27ac and H3K4me1 in primary ST cells, supporting its *cis*- regulatory identity (<u>fig. S7g</u>). Furthermore, Hi-C data confirmed higher-order chromatin interactions between the LTR8B element and the *STS* promoter in TSCs, which was consistent with the high expression of *STS* (<u>fig. S7g</u>). Taken together, our results indicate that a subset of LTR8B elements were co-opted as enhancers in regulating placental genes, and that SARS-CoV-2 infection led to the decreased activity of both the retrotransposon-derived enhancers and their target genes.

### Epigenetic dysregulation underlies decreased PSG expression in COVID-19 affected pregnancy

Given the transcriptional and chromatin state changes at the LTR8Bs and *PSGs*, we further aimed to delineate the underlying epigenetic mechanism. Integrating publicly available placental cell type RNA-seq datasets (*57, 58*), we confirmed that the *PSGs* are expressed at the highest levels in STs, followed by EVTs, and at relatively low levels in CTs, which are the undifferentiated trophoblasts (<u>fig. S8a</u>). Interestingly, in all analyzed cell types, the LTR8B elements within the *PSG* clusters were devoid of DNA methylation, suggesting that they were epigenetically regulated by alternative modifications (Fig. 7a). As such, we analyzed the enrichment of repressive histone marks on the LTR8Bs and found that H3K27me3, and not H3K9me3, was anti-correlated with the levels of H3K27ac (<u>fig. S8b, 8c</u>). These results suggest that the cell type-specific repression of these LTR8B elements was mediated by H3K27me3 and their dysregulation in SARS-CoV-2 infected pregnancies may be associated with aberrant epigenetic silencing.

It is also possible that the downregulation of *PSGs* is a result of reduced levels of TF binding, which normally target the LTR8B enhancers in STs. Motif analysis revealed that GATA was the most significantly enriched TF motif in LTR8B elements (Fig. 8c). Indeed, the GATA family of TFs are one of the master regulators in placenta development (*62*). Interestingly, we found that *GATA2* expression was significantly downregulated in ST and VCT clusters of patients (<u>fig. S8d</u>). Furthermore, we discovered that the target genes of GATA2 exhibited a general downregulated trend in patient ST cells (<u>fig. S8e</u>) (*63*). Taken together, we surmised that the LTR8B-derived enhancers, which were normally devoid of H3K27me3 in STs, regulate the *PSGs* via GATA2 binding. However, these retrotransposons showed reduced activity upon SARS-CoV-2 infection, which could be due to aberrant epigenetic repression and/or reduction of GATA2 recruitment.

## Discussion

The global COVID-19 pandemic, brought about by SARS-CoV-2 infection, impacted millions of people over the past two years. With the outbreaks of the current Delta and Omicron variants, hospitals have seen increasing numbers of SARS-CoV-2 infected pregnancies. Although mounting evidence suggests that COVID-19 is associated with an increased risk of adverse pregnancy outcomes, the precise molecular pathways underlying it remains unclear. In this study, the patients involved were diagnosed with COVID-19 at late pregnancy. Previous reports have suggested placental infection is rare but still possible (*2*). To discriminate the effect of maternal inflammatory response from a direct viral infection, we confirmed that the patient maternal-fetal interface samples were free from the virus at the time of analysis. Concordantly, we found low to undetectable expression of SARS-CoV-2 receptor *ACE2* and its cofactor *TMPRSS2* in our samples. This finding is consistent with that of a previous study showing that ACE2 and TMPRSS2 are expressed in a subpopulation of placental cells in early pregnancy and are rarely detected at term (*27*). Furthermore, we have observed a lower transfer ratio for SARS-CoV-2 specific IgG at 1.11 compared to >1.5 for other respiratory infections such as influenza or pertussis (Table S1) (*64*). Although three patients were IgG seropositive at delivery, only one neonate was IgG seropositive, which may be the result of the compromised placental transfer of anti-SARS-CoV-2 antibodies previously reported in cases with third trimester SARS-CoV-2 infection (*64*). We observed that even mild to asymptomatic SARS-CoV-2 infection in late pregnancy, without direct placental infection, was associated with immune and angiogenesis dysregulation at the maternal-fetal interface. Our findings suggest that these pregnancies could be at risk of placenta related complications and therefore specific care and management should be instigated for COVID-19 pregnant patients to rule out and monitor for preeclampsia and fetal growth restriction. Intriguingly, we found that retrotransposons, which have largely been neglected in prior studies, were dysregulated upon infection, providing a new direction for studying COVID-19 associated pregnancy complications.

The complex immunomodulation at the maternal-fetal interface plays an important role for successful pregnancies. Aberrant interferon expression is known to be a common cause of pregnancy disorders (*65*). Notably, several recent reports demonstrated immune activation and elevated interferon signalling at the maternal-fetal interface in SARS-CoV-2 infected pregnancies, particularly in T cells (*27–29*). Here, we detected upregulation of interferon-related genes across different immune cell types, including T cells, macrophages, dendritic cells, and NK cells, suggesting a more significant immune activation at the maternal-fetal interface than previously appreciated. This is further supported by the increased receptor-ligand interactions among immune cell clusters. Specifically, we found the concomitant upregulation of several IFI genes and IFITM genes. Importantly, experiments in mouse models have revealed that IFITM proteins inhibit Syncytin-mediated ST formation, which results in placental abnormalities (*66*). The process of CT fusing to form multi-nucleated ST occurs from the time of implantation until full term (*67*). Therefore, the overexpression of *IFITM* genes in late pregnancy can still potentially impair ST formation and function. A recent report suggests that the type I interferon response from SARS-CoV-2 infection can cause endothelial damage in the lungs by cGAS-STING activation (*68*), which is analogous to the placental endothelial damage observed in COVID-19 patients (*1, 4, 69, 70*). Taken together, we have found the aberrant expression of IFN-related genes across distinct immune cell types at the maternal-fetal interface, which is potentially detrimental to placental function and fetal health.

Apart from the increased risks of fetal demise and preterm birth, increased incidence of “preeclampsia-like syndrome” has been noted in pregnant patients with SARS-CoV-2 infection (*2*). Indeed, evidence suggests that preeclampsia and COVID-19 share similar clinical phenotypes of endothelial dysfunction and may have common pathophysiological mechanisms, especially those involving the renin-angiotensin system (RAS) (*70–72*). In healthy pregnancies, the RAS components contribute to invasion, migration, and angiogenesis of trophoblast cells (*73, 74*). Vasoconstriction, resulting from the suppression of RAS, has been demonstrated in both preeclampsia and SARS-CoV-2 infection (*5, 72, 75*). In conjunction with alteration of RAS, changes in expression and imbalance of angiogenic mediators, associated with SARS-CoV-2 infection, result in systemic endothelial dysfunction and the clinical syndrome of preeclampsia (*5, 47, 76*). Previous studies have shown endothelial damage, together with overexpression of proangiogenic factors, such as VEGF, basic fibroblast growth factor (FGF-2), and placental growth factors (PlGF) in the plasma or lung biopsies of COVID-19 patients (*77*). Furthermore, a recent study suggests a positive correlation between VEGF-A and PlGF levels with the severity of COVID-19 in nonpregnant patients (*78*). Abnormal angiogenesis, especially in the placenta, has been reported in pregnancy complications, however, the role of this mechanism in pregnant patients with COVID-19 needs further exploration (*44, 79*). We found that genes related to blood vessel formation were upregulated in endothelial cells and fibroblasts, concomitant with altered chromatin accessibility, which suggests the dysregulation of CREs. Furthermore, we found angiogenesis-related receptor-ligand pairs to have increased activity in patient samples, including *VEGF-FLT1* and *FGF-FGFR*. In particular, we discovered two putative enhancers upstream of *VEGF* gene, which gained chromatin accessibility along with the transcriptional upregulation of the target gene. In addition, we discovered the dysregulation of a novel angiogenesis gene *ENG*, which was upregulated and showed a concomitant change in chromatin states of its putative enhancers. Our findings support the involvement of such genes in endothelial dysfunction in the maternal-fetal interface of COVID-19 pregnant patients. Moreover, the epigenetic dysregulation of the related CREs could underlie their differential expression.

Although frequently neglected in genomic studies, increasing evidence shows retrotransposons to be integral parts of our genome, functioning as protein-coding genes and CREs (*15*). Indeed, specific retrotransposons play critical roles in placental development and function (*16–23*). It should also be noted that aberrant retrotransposon activities are associated with immune dysregulation in autoimmune diseases and cancers (*80–82*). Surprisingly, we found that hundreds of elements were differentially expressed and/or have altered chromatin states in patient samples, most of which were downregulated or had decreased chromatin accessibility. Such elements were associated with pregnancy genes and enriched with motifs for placenta development-related TFs, suggesting a role in placental functions. In particular, we discovered a subset of elements associated with the dysregulation of the *PSG* genes. *PSG* genes are immunoglobulin genes belonging to the carcinoembryonic antigen (CEA) family (*53*). The 10 *bona fide* human *PSG* genes cluster at chromosome 19q13 and contribute to the most abundant placenta-origin proteins in maternal blood, which are found to have immunoregulatory, pro-angiogenic, and anti-platelet functions (*53*). Copy-number variations and low levels of *PSGs* expression were reported to be associated with fetal growth restriction and preeclampsia (*54–56*). However, the regulation of these important placental factors remains largely unexplored. We have observed that the deregulation of LTR8B-derived enhancers is potentially responsible for the downregulation of *PSGs* in STs of COVID-19 patients. Our Hi-C data revealed the strong higher-order chromatin interactions between the retrotransposons and multiple *PSG* promoters, suggesting their contribution to the *PSG* regulatory modules. Incorporating public placental epigenomic datasets, we found that H3K27me3, rather than H3K9me3 or DNA methylation at these elements, was negatively correlated with the expression of *PSGs*, supporting the model that high *PSG* expression in normal STs is preceded by H3K27me3 reprogramming. In addition, these LTR8B elements are enriched with the GATA2 binding motif, and the downregulation of *PSG* coincides with the decreased expression of *GATA2*. We propose that aberrant H3K27me3 at retrotransposon-derived enhancers, coupled with downregulation of *GATA2,* is associated with decreased *PSG* expression in SARS-CoV-2 infection. These molecular changes potentially lead to impaired immunoregulation and angiogenesis and their interaction at the maternal-fetal interface and contribute to pregnancy complications in COVID-19 patients. It should be noted that the enhancer role of LTR8B elements was not restricted to the *PSG* cluster and another element was identified as a putative enhancer for the pregnancy-related *STS* gene. This subfamily’s impact may be a result of the conserved GATA2 binding motifs. Further investigation and functional validations are needed to better understand the importance of LTR8B elements in placental development and function.

Collectively, in this study, we generated extensive transcriptomic and epigenomic datasets of the maternal-fetal interface from SARS-CoV-2 infected patients and control pregnant patients, at both single-nucleus level and bulk levels. These serve as an important resource for the community to decipher the molecular alterations in COVID-19 pregnancies. Our findings reveal the critical role of epigenetic regulation in the COVID-19 pregnancy. We further defined novel retrotransposons-derived enhancers that are associated with the altered expression of important pregnancy genes. With the surge of the Omicron variant, there are vastly more cases of COVID-19 pregnancies, and further studies investigating whether the new variant could induce similar immune activation and angiogenesis dysregulation is needed. In summary, our study emphasizes the critical role of epigenetic regulation of non-coding elements in the maternal-fetal interface of pregnancies affected by COVID-19.

## Acknowledgments

We thank Qinghong Jiang and Dr Vincy Ho (HKUST) for their assistance in carrying out the study; Lijia Chen, Tracy CY Ma, Maggie Mak, and Angela ST Tai who were involved in study coordination; all participants and their attending obstetricians (Teresa Ma, Florrie NY Yu, Choi Wah Kong, Tsz Kin Lo, Po Lam So), nurses, midwives; and laboratory technicians at all participating hospitals (Queen Elizabeth Hospital, United Christian Hospital, Princess Margaret Hospital, Tuen Mun Hospital, Hong Kong, China) for case recruitment and sample collection. We also thank Prof Pengtao Liu for sharing the TSC cell line used in this study. Schematic was created with BioRender.com.

## Author contributions

L.G, V.M, S.K.M.T, X.Z, C.C.W, L.C.P, and D.L designed the study. G.E.G, S.M, and L.C.P collected all clinical samples and data. L.G, V.M, S.K.M.T, X.Z, L.Y.C, and B.W.L performed all experiments. L.G, V.M, S.K.M.T, X.Z, and M.F.C performed all data analyses. L.G, V.M, S.K.M.T, X.Z, C.C.W, L.C.P, and D.L prepared the manuscript.

## Data and materials availability

All sequencing datasets generated in this study have been deposited at ArrayExpress under the accession ID E-MTAB-11749 and at EGA under accession ID EGAS00001006263. Published trophoblasts epigenomic datasets were acquired from JGA under accession IDs JGA000074 and JGA000117. All other data are available in the main text or the supplementary materials.

## Supplementary Materials

Materials and Methods

Figs. S1 to S8

Tables S1 to S5

Data S1

