## Supplementary Materials for "SARS-CoV-2 impacts the transcriptome and epigenome at the maternal-fetal interface in pregnancy"

**This PDF file includes:**

Materials and Methods
Figs. S1 to S8
Tables S1 to S5

**Other Supplementary Materials for this manuscript include the following:**

Data S1. List of NGS datasets (.xlsx)

**Materials and Methods**

Patient consent and sample collection

This was a case-control study including consecutive pregnant patients who tested positive for SARS-CoV-2 infection by RT-qPCR of a deep throat saliva (DTS) or nasopharyngeal swab (NPS) sample, enrolled during pregnancy or at the time of delivery between 27 March 2020 and 24 January 2021. All participants gave written informed consent to participate in the study. In our study, seven patients with pregnancy beyond 24 weeks of gestation with positive RT-qPCR for SARS-CoV-2 during delivery were selected and the information regarding serial Ct values at and after diagnosis, demographic, clinical and outcome data and neonatal NPS/NPA RT-qPCR results were collected from medical records. Patients had blood drawn at the time of delivery and cord blood was collected immediately after delivery, and serum antibodies against SARS-CoV-2 were analyzed. As per hospital clinical protocols, SARS-CoV-2 infection was assessed using the cycle threshold (Ct) values obtained from the RT-qPCR assay applied to the DTS or NPS samples and the Ct value >35 was defined as negative. Qualitative detection of the anti-SARS-CoV-2 immunoglobulin G (IgG) directed against the nucleocapsid protein (N-protein) of the virus was carried out using the Elecsys Anti-SARS-CoV-2 assay (Roche, USA) on a cobas® e411 analyzer. The result was given either as cut-off index (COI) <1.0 (negative for anti-SARS-CoV-2 antibodies) or COI $^{3}$1.0 (positive for anti-SARS-CoV-2 antibodies). We also recruited seven SARS-CoV-2 uninfected women (as determined by negative RT-qPCR testing of DTS/NPS) who had undergone an elective Cesarean section and provided informed consent to donate the tissues to serve as uninfected controls for this study. Approval for the study was obtained from the Joint Chinese University of Hong Kong – New Territories East Cluster Clinical Research Ethics Committee (CREC Ref. No. 2020.210).

Maternal-fetal interface tissues were collected and processed immediately after delivery. All samples were grossly examined, and hematoxylin and eosin-stained sections were reviewed. Samples of the maternal-fetal interface including the decidua basalis and placental villi tissue were collected and snap freezing in liquid nitrogen for viral load detection or pulverized in liquid nitrogen into powder form for nuclei preparation and RNA extraction. Full thickness placental samples were fixed in formalin and paraffin embedded for histologic analyses according to standard methods.

Detection of SARS-CoV-2 in patient samples

Three pieces of tissue were used from each case and amplified in duplicate. Total RNA was extracted using RNeasy Mini Kit (QIAGEN). The detection of SARS-CoV-2 RNA was performed with the FDA-authorized CDC 2019-Novel Coronavirus (2019 nCoV) Real-Time RT-PCR Diagnostic Panel (EUA 200001). The nucleocapsid genes (both N1 and N2) were assayed, with the human RNase P (RP) as an endogenous reference control to verify that RNA was presented in every sample. Reactions (10006625, IDT) containing a DNA sequence of the SARS-CoV-2 N gene was used as positive controls. In addition, a set of *in vitro* synthesized RNA transcripts including 3 quantification positive controls (1000, 100 and 10 copies genome equivalent) were also assayed. No template control (NTC) wells were included as negative control.

Immunohistochemistry

Formalin-fixed paraffin embedded tissue blocks from all patients were cut into 5 μm thick sections for standard H&E and specific immunohistochemical (IHC) staining. The slides were deparaffinized with xylene and rehydrated using graded ethanol solution. For IHC, the slides were then treated with 3% hydrogen peroxide to deactivate the cellular peroxidases activities. Antigen retrieval was performed by heating the slides in a microwave oven using antigen retrieval buffer (pH 6.0) (Abcam, Cambridge, MA, USA). Then, incubated with Protein Block (Abcam, Cambridge, MA, USA) to block nonspecific background staining. The sections were then incubated overnight at 4°C with primary antibodies against FLNB (ab282106, Abcam), HDAC9 (ab52030, Abcam), NRP1 (ab81321, Abcam), PAPPA (ab174314 Abcam) and PSG9 (AP53483PU-N, Origene). After washing with 0.1% TBS- Tween-20 (TBS-T) to remove the unbound antibody, the sections were incubated with mouse anti-rabbit horseradish peroxidase (HRP)-conjugated secondary antibody (Sigma-Aldrich, St. Louis, MO, USA) for 30 minutes at room temperature. The slides were subsequently stained with DAB Substrate, 3,3'-diaminobenzidine, (EnVision, DAKO, Santa Clara, CA, USA) and counterstained with hematoxylin (Sigma-Aldrich, St. Louis, MO, USA).

Human trophoblast stem cell (TSC) culture

Human trophoblast stem cells (hTSCs) were a gift from Prof. Pengtao Liu, School of Biomedical Sciences, the University of Hong Kong *(59)*. TSC was cultured as previously described *(83)* with minor modifications. Briefly, the cells were maintained in TSC medium (DMEM/F12 supplemented with 0.1 mM 2-mercaptoethanol, 0.2% FBS, 0.5% Penicillin-Streptomycin, 0.3% BSA, 1% ITS-X supplement, 150µM L-ascorbic acid, 50 ng/ml EGF, 2 mM CHIR99021, 0.5 mM A83-01, 1 mM SB431542, 0.8 mM VPA and 5 mM Y27632). Tissue culture plates were coated with 5 µg/mL Collagen IV (Corning) at 37ºC for 1 hour. Cells were cultured at 37ºC, 5% $CO_2$. At approximately 80% confluency, cells were dissociated with TrypLE (Gibco) and passaged to new Collagen IV-coated plates. TSCs at 20-30 passages were harvested for RNA-seq, µChIP-seq, and Hi-C analyses.

Bulk RNA-seq and RT-qPCR

10-20 mg of pulverized tissue powder or 1-2 million TSCs pellet was homogenized in cold TRIzol reagent (Invitrogen), and RNA was extracted according to the manufacturer's manual. Extracted total RNA was used for generating bulk RNA-seq libraries and for RT-qPCR. For bulk RNA-seq, 1 µg total RNA was used as input. Briefly, ribosomal RNA (rRNA) was removed using Ribo-off rRNA Depletion Kit (Vazyme). The rRNA-depleted total RNA was subsequently proceeded to RNA-seq library preparation using QIAseq Stranded Total RNA Library Kit (QIAGEN) as described in the manufacturer's manual. The RNA-seq libraries were sequenced on Illumina Nextseq 500 platform.

For RT-qPCR analyses of dysregulated genes and retrotransposons, which were carried out separately from the SARS-CoV-2 diagnostic RT-qPCR, 1 µg total RNA was treated with DNase I (NEB) and purified with RNAClean XP beads (Beckman Coulter) before first-strand synthesis. First-strand synthesis was carried out using Superscript III Reverse Transcription System (Thermo Fisher Scientific) according to the manufacturer's manual. cDNA was then analyzed by qPCR on LightCycler 480 Instrument II. Primers used in this study are listed in Supplementary Table 5.

Bulk ATAC-seq library preparation

Bulk ATAC-seq protocol was adopted from *(84)* with minor modifications. Briefly, 10-20 mg pulverized tissue powder was transferred to a pre-chilled 1.5ml LoBind tube (Eppendorf) and immediately resuspended in 1 mL nuclei permeabilization buffer (5% BSA, 0.2% NP-40, 1mM DTT, 1× protease inhibitors in PBS) and rotated for 10 minutes at 4°C. The nuclei suspension was filtered through a 40 μm Cell Strainer (Corning) to remove large clumps of tissues. Filtered nuclei were spun down at 500 × g, 4°C for 5 minutes, and resuspended in 50 μl of chilled tagmentation buffer (10 μL 5 × TTBL buffer (Vazyme), diluted in nuclease-free water). An aliquot of the suspension was taken for counting by hemocytometer and the nuclei concentration was then adjusted to 2,000 – 5,000 nuclei/μL. 0.5 μL Vazyme V50 Tn5 transposase was added to 9.5μl nuclei suspension and the reaction was incubated in a thermomixer at 37°C with 500 rpm mixing for 30 minutes. The tagged DNA was amplified with KAPA HiFi Hotstart Ready Mix (Roche) for 5-10 PCR cycles, followed by size selection with AmpureXP beads (Beckman Coulter). The libraries were sequenced on Illumina Nextseq 500 platform.

snRNA-seq and snATAC-seq library preparation

snRNA-seq and snATAC-seq were conducted according to the manufacturer's protocols with minor modifications. 10-20 mg pulverized tissue powder was transferred to a pre-chilled 1.5 ml LoBind tube (Eppendorf) and immediately resuspended in 1 mL nuclei permeabilization buffer (5% BSA, 0.2% NP-40, 1mM DTT, 1× protease inhibitors in PBS, 40U/μl RNase inhibitor) and rotate for 10 minutes at 4°C. The nuclei suspension was filtered through a 40 μm Cell Strainer (Corning) to remove clumps of tissues. The nuclei were spun down at 500 × g, 4°C for 5 minutes and then ran on the Chromium Next GEM Single Cell 3′ V3.1 platform for snRNA-seq, and on the Chromium Next GEM Single Cell ATAC v1.1 platform for snATAC-seq. Libraries from 10x Genomics platforms were then converted for sequencing on the MGI sequencing platform with MGIEasy Universal Library Conversion Kit. Converted libraries were sequenced on the MGISEQ-2000RS platform.

Hi-C library preparation

TSC Hi-C library was generated with Arima-Hi-C Kit according to the manufacturer's protocol. The Hi-C library was converted for sequencing on the MGI sequencing platform with the MGIEasy Universal Library Conversion Kit. The converted library was sequenced on MGISEQ-2000RS platform.

μChIP-seq library preparation

TSC μChIP-seq libraries were prepared as described previously *(85)* with minor modifications. Briefly, $5 \times 10^5$ crosslinked cells were resuspended in lysis buffer and sonicated with the Covaris S220 sonicator continuously for 400s at 175W. Fragmented chromatin was added to Protein A Dynabeads (Thermo Fisher Scientific) that were bound beforehand to antibodies against H3K27ac (AM39133, Active Motif), H3K4me3 (AM39915, Active Motif), and H3K4me1 (AM91289, Active Motif). The mixtures were then incubated at 4 °C for 40 hours with rotation. Captured chromatin was washed four times with RIPA buffer and subsequently eluted at 37 °C for 1 hour in elution buffer. Reverse crosslinking was performed by incubation at 68 °C for 4 hours with Proteinase K. Eluted DNA was purified with the QIAquick PCR Purification Kit (QIAGEN). Libraries were prepared using the KAPA HyperPrep Kit (Roche) according to the manufacturer's protocol and sequenced on Illumina Nextseq 500 platform.

Luciferase assay

Dual-Luciferase® Reporter Assay System (Promega E1960) was used to perform luciferase assay for *PSG* intronic LTR8B elements of interest in TSC cells. Briefly, the LTR8B elements of interest or GFP sequence (background negative control) were cloned into the pGL3-promoter and pGL3-enhancer vectors. The primers used for amplification of target sequences are listed in Supplementary Table 5. For transfection, TSCs were seeded at $4 \times 10^4$ per well in 24-well plates 2 days prior to transfection. Cells were then co-transfected with the *Renilla* luciferase vector and pGL3 vector containing an LTR8B element or GFP sequence with Lipofectamine 3000. Luciferase activity was measured using Dual-Luciferase Reporter Assay System reagents (Promega E1960) with Spectronic Genesys 5 UV/ Visible Spectrophotometer (ALT) 48 hours after transfection. Luciferase activity was normalized to both *Renilla* luciferase signal and pGL3 basic vector signal.

Bulk RNA-seq data analysis

RNA-seq datasets from maternal-fetal interface samples, TSCs, and publicly available human placenta datasets were analyzed according to the following process. Reads were aligned to the GRCh38/hg38 genome assembly and the GECODE V39 transcriptome assembly separately with STAR v2.5.3a *(86)* with the parameters --outFilterMultimapNmax 1 --alignSJoverhangMin 8 --alignSJDBoverhangMin 1 --outFilterMismatchNmax 999 --outFilterMismatchNoverReadLmax 0.04 --alignIntronMin 20 --alignIntronMax 1000000 --alignMatesGapMax 1000000 --outSAMstrandField intronMotif --quantMode TranscriptomeSAM --sjdbScore 1. Only uniquely mapped reads were kept in both alignments. Transcriptome alignments were then quantified by RSEM *(87)* with parameters --bam --estimate-rspd --calc-ci --seed 12345 --no-bam-output --ci-memory 70000 --paired-end to calculate fragments per kilobase per million reads (FPKM) values of annotated transcripts. Genomic alignments were piled up to generate reads per million (RPM) signals and used for visualization. For retrotransposons quantification, coordinates for retrotransposons were assigned based on RepearMasker annotations. Paired-end libraries were treated as single-end and uniquely aligned reads were used to quantify individual elements' expression. Differentially expressed genes and retrotransposons were defined using DESeq2 v1.22.1 *(88)* with default settings.

Bulk ATAC-seq data analysis

Bulk ATA-seq datasets from maternal-fetal interface samples were analyzed according to the following process. Reads with low quality (Phred score < 20) were removed and adaptors were trimmed by trim_galore v0.4.3 under paired-end mode. Reads were then aligned to GRCh38/hg38 genome assembly using Bowtie2 v2.3.3.1 *(89)* with parameters -N 1 -L 25 -X 2000 --no-discordant --no-mixed. Duplicated uniquely aligned reads were further removed by Picard MarkDuplicates v2.9.0. To make the start site of each read represent the centre of the transposase binding *(90)*, the alignments were adjusted using MACS2 v2.1.0 *(91)* with parameters –shift -75 –extsize 150.

To normalize the ATAC-seq signal and background, S3norm was applied *(92)*. In brief, the whole-genome coverage (20bp bin) was calculated from raw read counts using "bigWigAverageOverBed". The converted files together with background (coverage of all bins = 1) were then normalized by S3norm with the default setting. RPM values were then generated for visualization. Peaks were called by MACS2 with parameters -c 2 -l 100 -p 0.01 using the S3norm output. Peaks called from all samples were merged and differential peaks were defined by DESeq2 with the S3norm normalized read counts. Differential retrotransposon chromatin accessibility was assigned by overlapping the retrotransposons coordinate with the defined differential peaks.

snRNA-seq data analysis

snRNA-seq raw fastq files generated from the MGISEQ-2000RS platform were converted to 10x Cell Ranger compatible format. The converted fastq files were mapped by cellranger v3 mapped to GRCh38 pre-mRNA genome and the SARS-CoV-2 genome (severe acute respiratory syndrome coronavirus 2 isolate Wuhan-Hu-1, GenBank NC_045512.2) with default setting. The output was then filtered and processed using Seurat v3 *(93)*. We used standard filtering to call single cells (nUMI ≥ 1000, nGene ≥ 1000, nUMI < 25000, nGene < 4000 and mitochondria ratio < 0.05) and removed clusters with a high droplet score, and no marker gene expression. Cells that passed the filtering (n=37,096) were normalized and transformed using Seurat's SCTransform function with the glmGamPoi method. The cells were then subclustered and annotated based on manual curation of markers, and reference to placenta marker databases *(31-33)*.

Each cell type from patient and control samples were analyzed for differentially expressed genes using Seurat's FindMarker function (p-adjusted value < 0.0.5, avg_log2(FoldChange) > 0.25) for each cluster. CellPhoneDB v3 *(42)* was used to analyze receptor-ligand interactions using standard processing and subsampling of 5000 cells was done for both patient and control samples. The receptor-ligand pairs were filtered for significant pairs only present in patient samples.

Pseudo-bulk signals of each cell type from patient or control samples were generated according to the following processes. In brief, raw reads of individual nuclei were split into separate files based on cell barcode. Reads from each nucleus were aligned to GRCh38/hg38 genome assembly with STAR v2.5.3a as described in bulk RNA-seq processing and only uniquely mapped reads were kept. Aligned reads with the same UMI were further removed by UMI_Tools v0.2.3 dedup function *(94)*. Genomic alignments of all nuclei under the same cell type from patient or control samples were merged to generate the pseudo-bulk RPM signals and used for visualization. For retrotransposons quantification, coordinates for retrotransposons were assigned based on RepeatMasker annotation and uniquely aligned reads were used to quantify signals from individual retrotransposon elements.

snATAC-seq data analysis

snATAC-seq raw fastq files generated from MGISEQ-2000RS platform were converted to 10x Cell Ranger ATAC compatible format. Converted fastq files were mapped by cellranger-atac v2.0.0 to GRCh38/hg38 assembly with default setting. Mapped snATAC-seq files were filtered and annotated with ArchR v1.0.1 *(34)*. Briefly, to filter for high quality nuclei, only nuclei with minimum number of fragments of 1,000, minimum TSS signal of 4, and minimum promoter ratio of 0.075 were kept. After initial clustering, we removed clusters with low overall TSS signals, which yielded 12,191 control nuclei and 11,710 patient nuclei (total N = 23,901). To annotate the cell types, we performed label transfer from snRNA-seq dataset with ArchR.

For peak analysis, pseudo-bulk signals for each cell type from patient or control samples were generated according to the following processes. In brief, raw reads of individual nuclei were split into separate files based on their cell barcodes. Duplicated reads within each nucleus were removed by FastUniq v1.1 *(95)*. Reads from each nucleus were aligned to GRCh38/hg38 assembly by STAR v2.5.3a and only uniquely aligned reads were retained. To make the start site of each read represent the centre of the transposase binding, the alignments were adjusted using MACS2 v2.2.7.1 with parameters –shift -75 –extsize 150. Genomic alignments of the nuclei of the same cell type were subsequently merged to generate pseudo-bulk signals for peak calling and visualization. Peak calling was performed using MACS2, with parameters –nomodel –keep-dup all -q 0.01 to define open chromatin sites. Peaks called from each cluster were merged to generate a master peak set, which was used for Peak2GeneLinkage analysis with ArchR to define potential *cis*-regulatory enhancers, and differentially accessible peak analysis. To define cell type-specific differentially accessible peaks in patients versus controls, we conducted Poisson test on normalized pseudo-bulk reads count, and significantly differentially accessible peaks were defined by fold change > 2, adjusted P-value < 0.1, and presence of signal in more than 5% of cells in the corresponding cell cluster. Differentially accessible individual retrotransposons were defined by those elements that overlapped with differentially accessible peaks.

Retrotransposon subfamily analysis

We used our Subfamily Assignment for Multiple Alignment (SAMA) pipeline *(51)* for retrotransposon subfamily quantifications in bulk and single-nucleus RNA-seq and ATAC-seq. Briefly, reads from each nucleus for single-nucleus data or from each individual sample for bulk data were mapped to GRCh38/hg38 assembly using STAR v2.5.3a with the parameter --outFilterMultimapNmax 150, and reads with more than one best genomic alignment that is uniquely anchored to the same repeat subfamily were rescued. Subsequently, rescued multi-aligned reads and uniquely aligned reads were integrated to quantify the expression or chromatin accessibility signal for each retrotransposon subfamily. To define differentially expressed or accessible retrotransposon subfamilies, fold change was calculated from the average signal of each subfamily across samples for bulk RNA-seq and ATAC-seq or for individual nuclei of the same cell type for snRNA-seq and snATAC-seq. P-value was calculated by Student's T-test.

Motif, Gene Ontology and retrotransposons enrichment analysis

Motif analysis was conducted by HOMER *(96)* with the whole genome as background. Gene Ontology analysis for differentially expressed genes and retrotransposons was generated using Metascape v3.5 *(97)* and GREAT v4.0.4 *(98)*, respectively. The top significant terms were shown. Enrichment of differentially expressed individual retrotransposons in a subfamily was calculated by the ratio of observed over expected counts of elements in each subfamily. The expected numbers were estimated by (n/N)*X (n=total number of elements of each subfamily; N=total number of retrotransposons in the genome; X=total number of differential retrotransposons). P-value was calculated by hypergeometric test.

ChIP-seq data analysis

For TSC μChIP-seq (single-end) datasets generated in this study, reads were aligned to GRCh38/hg38 assembly by Bowtie v1.3.0 *(99)* with parameters -v 3 - m 1 --best --strata, keeping uniquely aligned reads with no more than 3 mismatches, and no more than one best alignment was kept, hence no additional steps were needed to remove multi-aligned reads. PCR duplicates were removed by Picard MarkDuplicates v2.23.4.

For public placenta ChIP-seq datasets obtained from the IHEC data repository, paired-end reads were aligned to GRCh38/hg38 assembly by Bowtie2 v7.5.0 with parameters -N 1 -L 25 -X 500 --no-discordant --no-mixed. Reads with more than one best alignment were removed. PCR duplicates were removed by Picard MarkDuplicates v2.23.4. For quantification of H3K27me3 and

H3K9me3 enrichment at the LTR8B elements, input subtracted RPKM values were calculated for each element.

Hi-C data analysis

TSC Hi-C reads were mapped with Juicer v1.13 *(100)* to GRCh38/hg38 assembly with parameter -s Arima. All other parameters were set to default. Significant interactions were called by Fit-Hi-C at 5kb resolution, and interactions with $q < 0.05$ were kept. For virtual 4C, we extracted the reads that interacted with the bait (LTR8B elements of interest) and their flanking 2kb regions. The extracted reads were subsequently aligned to the genome with BWA v0.7.15 with parameter mem, and the RPKM signal was calculated for visualization.

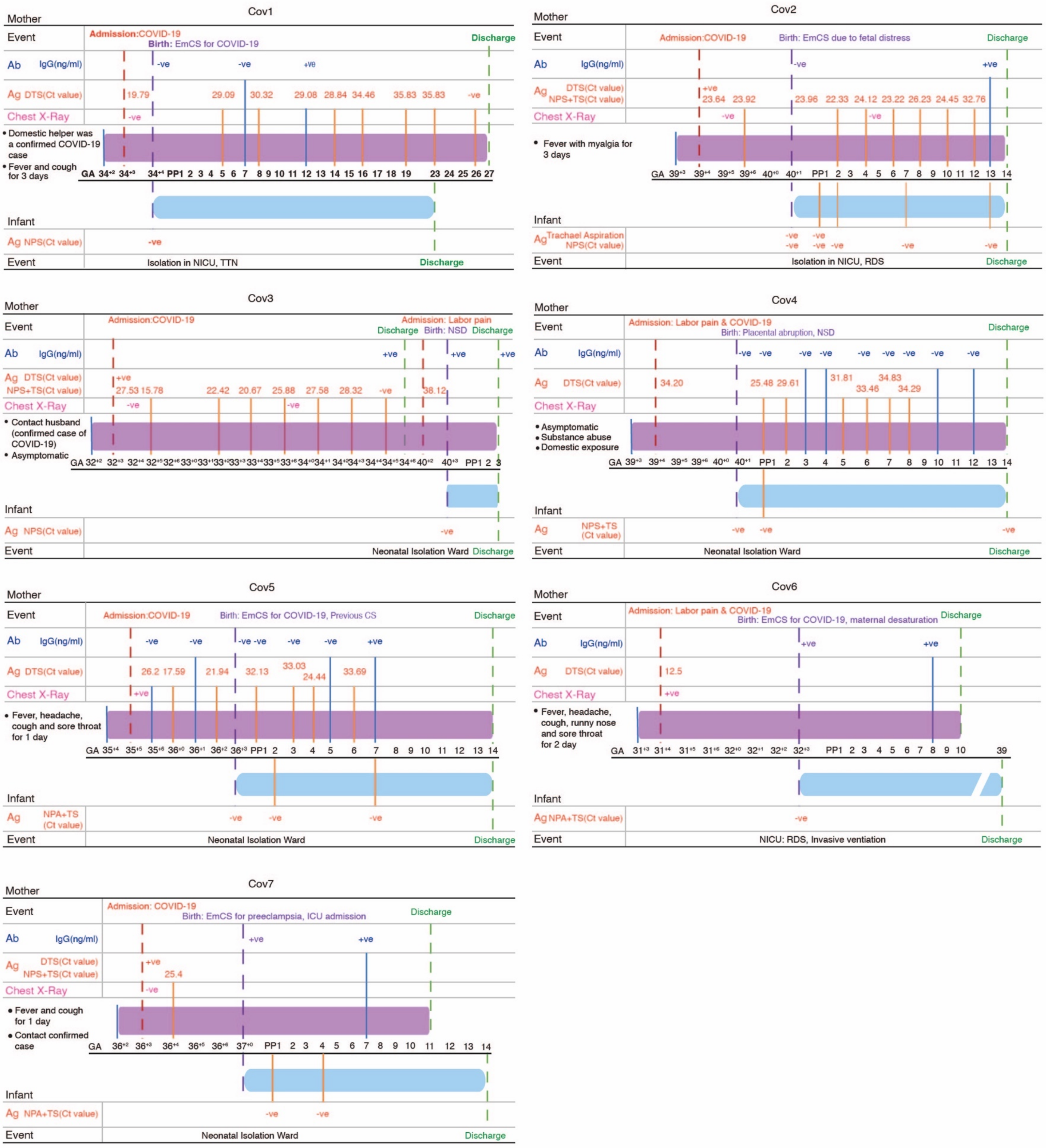
Cov1
Mother
Event
Admission:COVID-19
Birth: EmCS for COVID-19
Discharge
Ab IgG(ng/ml)
-ve -ve +ve
Ag DTS(Ct value)
19.79 29.09 30.32 29.08 28.84 34.46 35.83 35.83 -ve
Chest X-Ray
-ve
• Domestic helper was a confirmed COVID-19 case
• Fever and cough for 3 days
GA 34+2 34+3 34+4 PP1 2 3 4 5 6 7 8 9 10 11 12 13 14 15 16 17 18 19 23 24 25 26 27
Infant
Ag NPS(Ct value)
-ve
Event
Isolation in NICU, TTN
Discharge
Cov2
Mother
Event
Admission:COVID-19
Birth: EmCS due to fetal distress
Discharge
Ab IgG(ng/ml)
-ve +ve
Ag DTS(Ct value) NPS+TS(Ct value)
+ve 23.64 23.92 23.96 22.33 24.12 23.22 26.23 24.45 32.76
Chest X-Ray
-ve -ve
• Fever with myalgia for 3 days
GA 39+3 39+4 39+5 39+6 40+0 40+1 PP1 2 3 4 5 6 7 8 9 10 11 12 13 14
Infant
Ag Trachael Aspiration NPS(Ct value)
-ve -ve -ve -ve -ve -ve -ve -ve
Event
Isolation in NICU, RDS
Discharge
Cov3
Mother
Event
Admission:COVID-19
Discharge
Admission: Labor pain
Birth: NSD
Discharge
Ab IgG(ng/ml)
+ve +ve +ve
Ag DTS(Ct value) NPS+TS(Ct value)
+ve 27.53 15.78 22.42 20.67 25.88 27.58 28.32 -ve 38.12
Chest X-Ray
-ve -ve
• Contact husband (confirmed case of COVID-19)
• Asymptomatic
GA 32+2 32+3 32+4 32+5 32+6 33+0 33+1 33+2 33+3 33+4 33+5 33+6 34+0 34+1 34+2 34+3 34+4 34+5 34+6 40+2 40+3 PP1 2 3
Infant
Ag NPS(Ct value)
-ve
Event
Neonatal Isolation Ward
Discharge
Cov4
Mother
Event
Admission: Labor pain & COVID-19
Birth: Placental abruption, NSD
Discharge
Ab IgG(ng/ml)
-ve -ve -ve -ve -ve -ve -ve -ve -ve
Ag DTS(Ct value)
34.20 25.48 29.61 31.81 33.46 34.83 34.29
Chest X-Ray
• Asymptomatic
• Substance abuse
• Domestic exposure
GA 39+3 39+4 39+5 39+6 40+0 40+1 PP1 2 3 4 5 6 7 8 9 10 11 12 13 14
Infant
Ag NPS+TS (Ct value)
-ve -ve -ve
Event
Neonatal Isolation Ward
Discharge
Cov5
Mother
Event
Admission:COVID-19
Birth: EmCS for COVID-19, Previous CS
Discharge
Ab IgG(ng/ml)
-ve -ve -ve -ve -ve -ve +ve
Ag DTS(Ct value)
26.2 17.59 21.94 32.13 33.03 24.44 33.69
Chest X-Ray
+ve
• Fever, headache, cough and sore throat for 1 day
GA 35+4 35+5 35+6 36+0 36+1 36+2 36+3 PP1 2 3 4 5 6 7 8 9 10 11 12 13 14
Infant
Ag NPA+TS (Ct value)
-ve -ve -ve
Event
Neonatal Isolation Ward
Discharge
Cov6
Mother
Event
Admission: Labor pain & COVID-19
Birth: EmCS for COVID-19, maternal desaturation
Discharge
Ab IgG(ng/ml)
+ve +ve
Ag DTS(Ct value)
12.5
Chest X-Ray
+ve
• Fever, headache, cough, runny nose and sore throat for 2 day
GA 31+3 31+4 31+5 31+6 32+0 32+1 32+2 32+3 PP1 2 3 4 5 6 7 8 9 10 39
Infant
Ag NPA+TS(Ct value)
-ve
Event
NICU: RDS, Invasive ventiation
Discharge
Cov7
Mother
Event
Admission: COVID-19
Birth: EmCS for preeclampsia, ICU admission
Discharge
Ab IgG(ng/ml)
+ve +ve
Ag DTS(Ct value) NPS+TS(Ct value)
+ve 25.4
Chest X-Ray
-ve
• Fever and cough for 1 day
• Contact confirmed case
GA 36+2 36+3 36+4 36+5 36+6 37+0 PP1 2 3 4 5 6 7 8 9 10 11 12 13 14
Infant
Ag NPA+TS(Ct value)
-ve -ve
Event
Neonatal Isolation Ward
Discharge

**Fig. S1. Case timelines for the Patient samples.** The timeline of SAR-CoV-2 infection for each COVID-19 patient participating in this study. The timelines show the relationship between infection period, hospital admission due to SAR-CoV-2 infection, hospital admission for delivery and postpartum period. COVID-19 = Coronavirus disease 2019; Ab = Antibody; IgG = Immunoglobulin G; Ag = Antigen; DTS = Deep throat saliva; NPS = Nasopharyngeal swab; NPA = Nasopharyngeal aspiration; TS = Throat swab; CS = Cesarean section; EmCS = Emergency Caesarean section; RDS = Respiratory distress syndrome; Ct = Cycle threshold; NSD = Normal spontaneous delivery; TTN = Transient tachypnea of the newborn; ICU = Intensive Care Unit; NICU = Neonatal Intensive Care Unit; +ve = Positive finding; -ve = Negative finding; PP = Postpartum; GA = Gestational age.

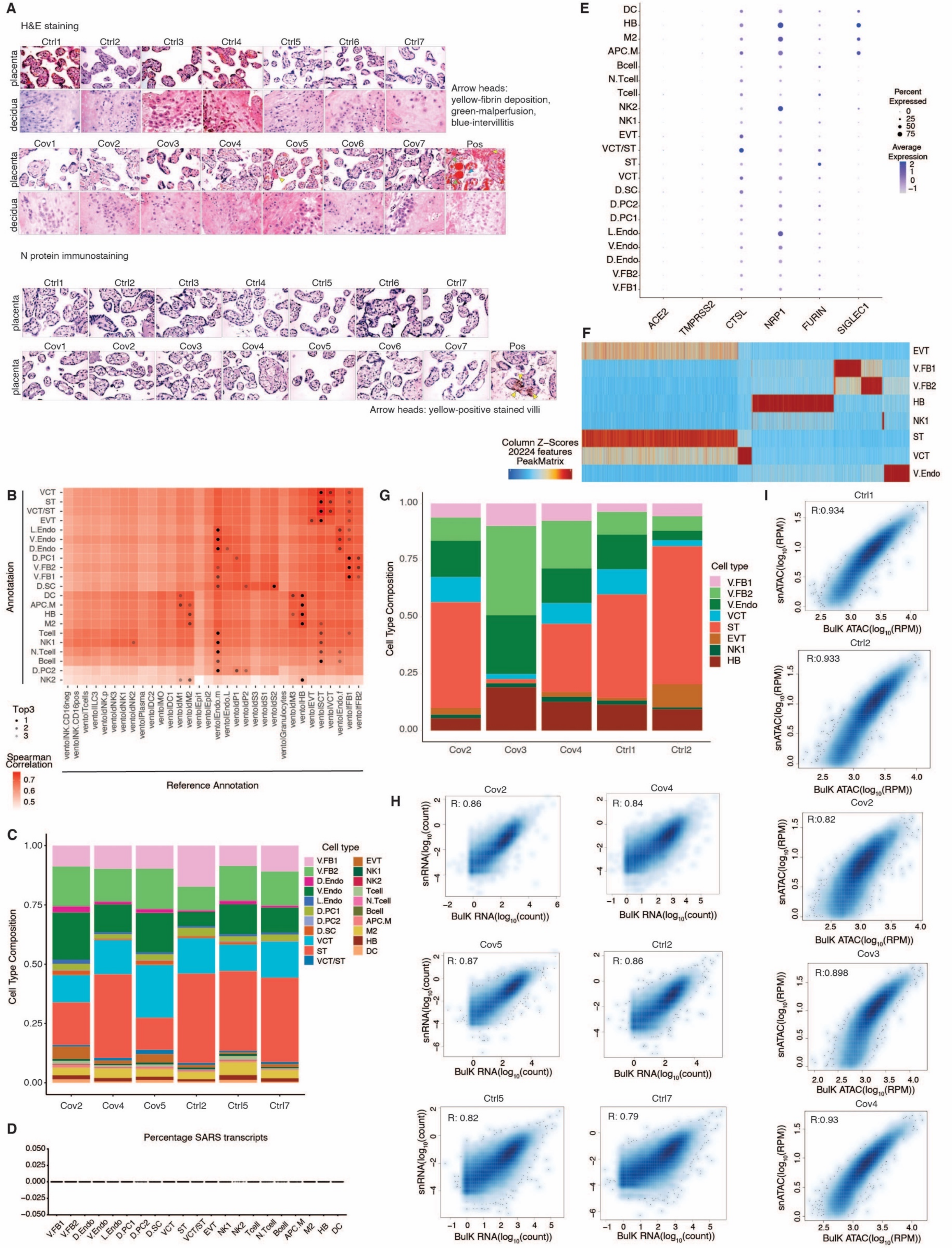
A
H&E staining
Ctrl1
Ctrl2
Ctrl3
Ctrl4
Ctrl5
Ctrl6
Ctrl7
placenta
decidua
Arrow heads:
yellow-fibrin deposition,
green-malperfusion,
blue-intervillitis
Cov1
Cov2
Cov3
Cov4
Cov5
Cov6
Cov7
Pos
N protein immunostaining
Arrow heads: yellow-positive stained villi
B
Annotation
Reference Annotation
Top3
Spearman Correlation
C
Cell Type Composition
Cell type
Cov2
Cov4
Cov5
Ctrl2
Ctrl5
Ctrl7
D
Percentage SARS transcripts
E
ACE2
TMPRSS2
CTSL
NRP1
FURIN
SIGLEC1
Percent Expressed
Average Expression
F
Column Z-Scores
20224 features
PeakMatrix
G
Cell Type Composition
Cell type
Cov2
Cov3
Cov4
Ctrl1
Ctrl2
H
snRNA(log10(count))
BulK RNA(log10(count))
R: 0.86
R: 0.84
R: 0.87
R: 0.86
R: 0.82
R: 0.79
I
snATAC(log10(RPM))
BulK ATAC(log10(RPM))
R:0.934
R:0.933
R:0.82
R:0.898
R:0.93

**Fig. S2**. **Quality control for multi-omic datasets, details on analyzed cell types, and validation of patient samples. (A)** Hematoxylin and eosin (H&E) (top) histological staining of placenta and decidua tissues from patients and controls. Yellow arrowheads indicate fibrin deposition, green arrowheads indicate malperfusion and blue arrowheads indicate intervillitis. Immunostaining of SARS-CoV-2 Nucleocapsid protein (bottom) in patient and control placenta tissues. Yellow arrowheads indicate positive stained villi. **(B)** Heatmap comparing our cell type annotation and public annotation (*32*). The color indicates the Spearman correlation and the black dot indicates the top 3 genes. **(C)** Stacked bar chart showing cell number and cell type distribution between Patient and Control samples in snRNA-seq after filtering. The color corresponds to annotated distinct cell types from the snRNA-seq. **(D)** Violin plots showing ratio of transcripts mapped to SARS-CoV-2 transcriptome in each cell type in snRNA-seq. Note that almost all points reside at 0. **(E)** Bubble plot showing the expression patterns of known SARS-CoV-2 receptors in each cell type, as measured by snRNA-seq. The size of the bubble indicates the percentage of cells within the given cluster that show expression and color indicates the average expression within the cluster. **(F)** Heatmap of snATAC-seq defined peaks on marker genes for each cell type. The color indicates the z-score of chromatin accessibility. **(G)** Stacked bar chart showing cell number and cell type distribution between all samples in snATAC-seq after filtering. The color corresponds to annotated distinct cell types in the snATAC-seq. **(H)** Scatter plots showing the Pearson correlation between the gene counts in snRNA-seq and Bulk RNA-seq. Genes were filtered for those have counts in either snRNA-seq or bulk RNA-seq. **(I)** Scatter plots showing Pearson correlation between the signal as measured by snATAC-seq and bulk ATAC-seq at transcriptional start sites (TSS). TSSs were filtered for those with signal in either snATAC-seq or bulk ATAC-seq.

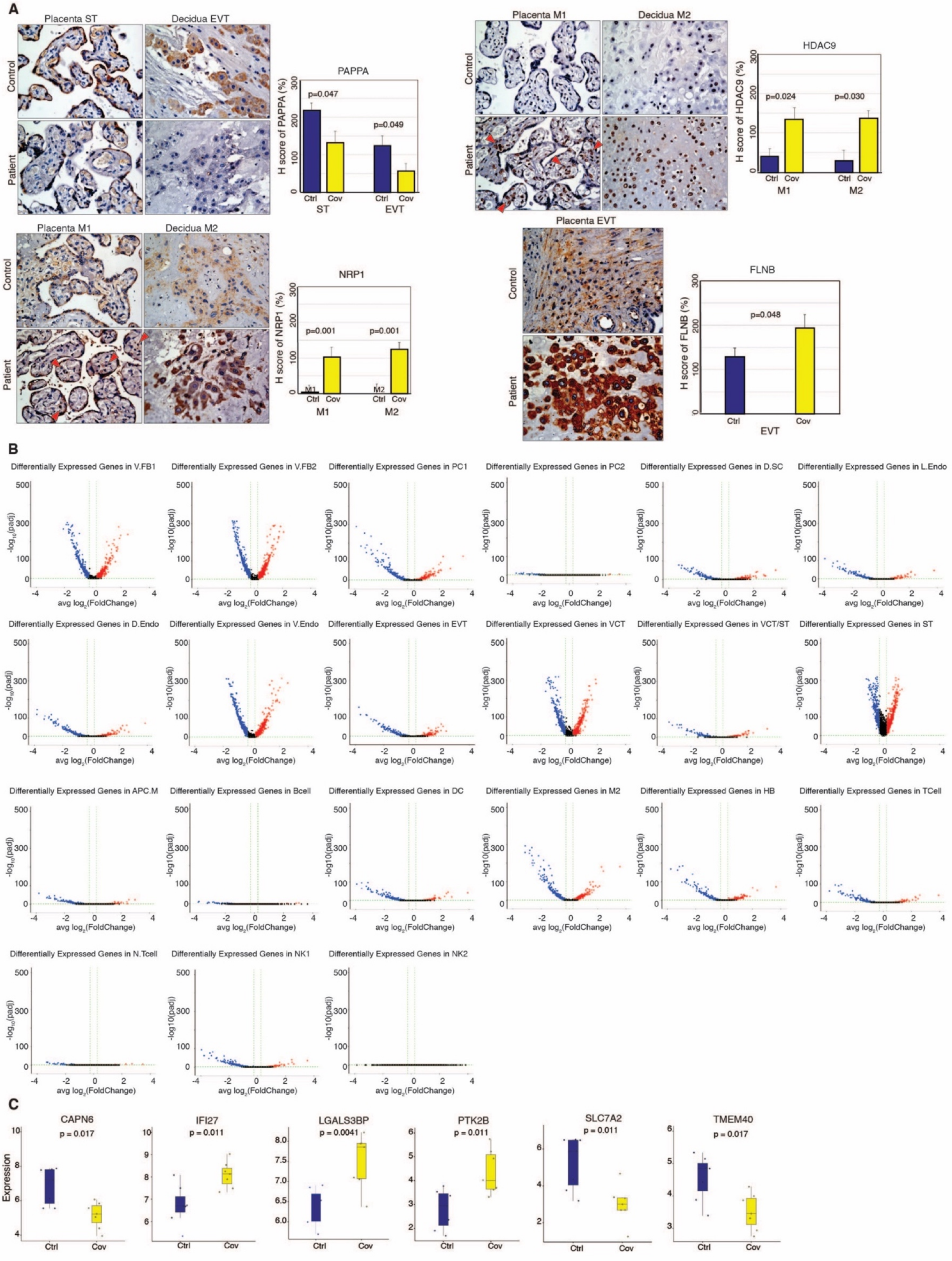
A
Placenta ST
Decidua EVT
Control
Patient
PAPPA
H score of PAPPA (%)
p=0.047
p=0.049
Ctrl
Cov
ST
EVT
Placenta M1
Decidua M2
HDAC9
H score of HDAC9 (%)
p=0.024
p=0.030
M1
M2
NRP1
H score of NRP1 (%)
p=0.001
p=0.001
Placenta EVT
FLNB
H score of FLNB (%)
p=0.048
B
Differentially Expressed Genes in V.FB1
Differentially Expressed Genes in V.FB2
Differentially Expressed Genes in PC1
Differentially Expressed Genes in PC2
Differentially Expressed Genes in D.SC
Differentially Expressed Genes in L.Endo
Differentially Expressed Genes in D.Endo
Differentially Expressed Genes in V.Endo
Differentially Expressed Genes in EVT
Differentially Expressed Genes in VCT
Differentially Expressed Genes in VCT/ST
Differentially Expressed Genes in ST
Differentially Expressed Genes in APC.M
Differentially Expressed Genes in Bcell
Differentially Expressed Genes in DC
Differentially Expressed Genes in M2
Differentially Expressed Genes in HB
Differentially Expressed Genes in TCell
Differentially Expressed Genes in N.Tcell
Differentially Expressed Genes in NK1
Differentially Expressed Genes in NK2
-log10(padj)
avg log2(FoldChange)
C
CAPN6
p = 0.017
IFI27
p = 0.011
LGALS3BP
p = 0.0041
PTK2B
p = 0.011
SLC7A2
p = 0.011
TMEM40
p = 0.017
Expression
Ctrl
Cov

D

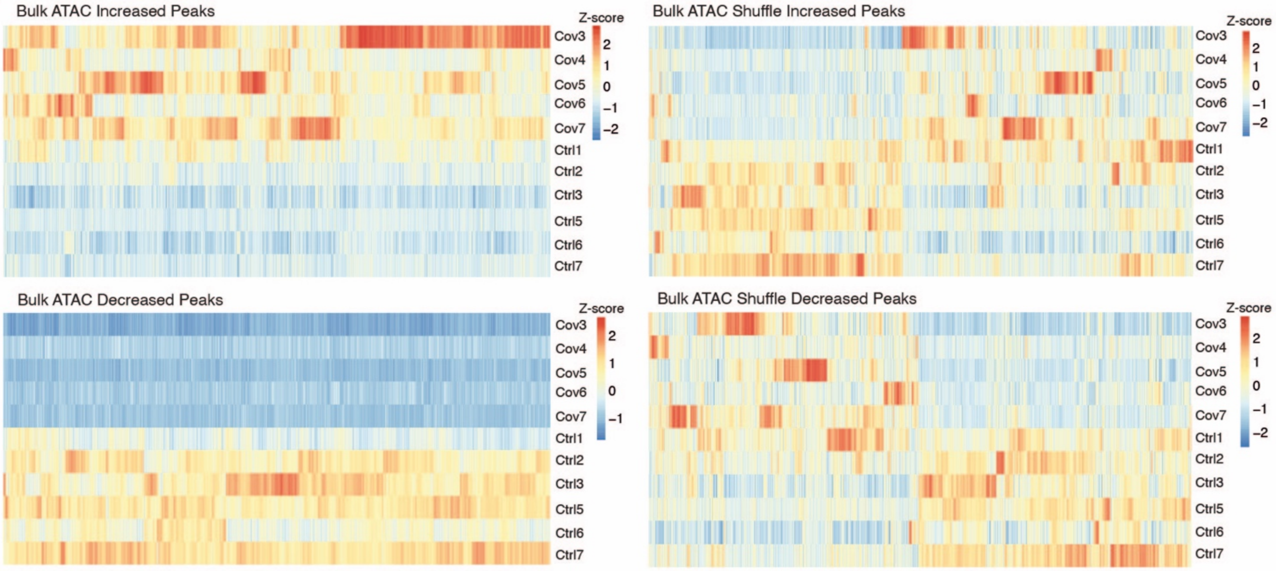


E

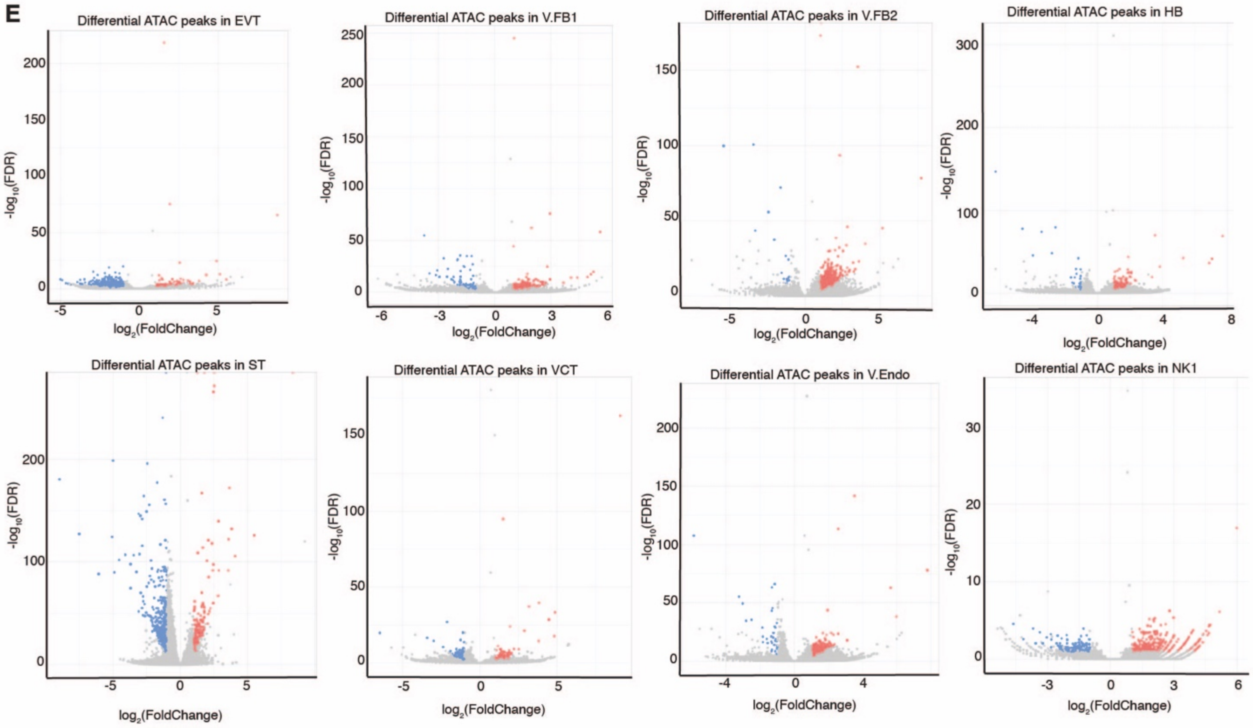


F

Decreased Peaks MOTIFs

| Motif | Transcription factor | P-value | # Sequence with motif |
|---|---|---|---|
| 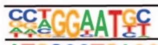 | TEAD4 | 1e-580 | 2937 |
| 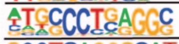 | AP-2α | 1e-504 | 2139 |
| 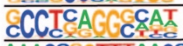 | AP-2γ | 1e-502 | 2476 |
| 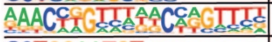 | GRHL2 | 1e-492 | 1683 |
| 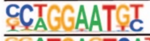 | TEAD2 | 1e-436 | 2083 |
| 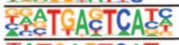 | FOLS1 | 1e-307 | 1723 |
| 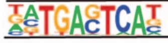 | BATF | 1e-294 | 1912 |

Increased Peaks MOTIFs

| Motif | Transcription factor | P-value | # Sequence with motif |
|---|---|---|---|
| 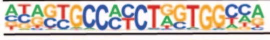 | CTCF | 1e-458 | 952 |
| 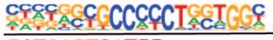 | CTCFL | 1e-315 | 1141 |
| 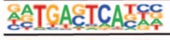 | FOSL2 | 1e-118 | 621 |
| 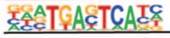 | FOSL1 | 1e-113 | 781 |
| 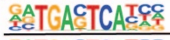 | JUN-AP1 | 1e-110 | 503 |
| 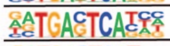 | ATF3 | 1e-104 | 866 |
| 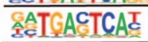 | JUNB | 1e-101 | 776 |

G

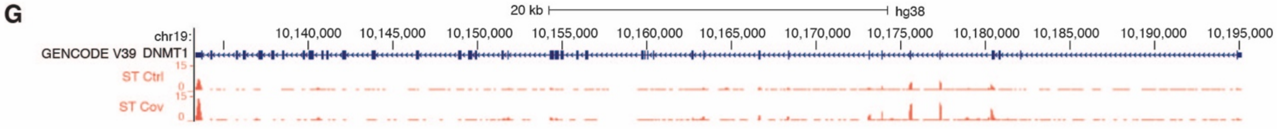

**Fig. S3. Characterization of transcriptional and chromatin accessibility changes in patient samples. (A)** Immunostaining and corresponding H-score as a percentage (right) of differentially expressed genes PAPPA, NRP1, HDAC9, and FLNB respectively in placenta and decidua tissue. **(B)** Volcano plots showing dysregulated genes in all 21 cell types as defined from snRNA-seq. Upregulated genes (red) (padj <0.05 and $log_2$(fold change) > 0.25) and downregulated genes (blue) (padj <0.05 and $log_2$ (fold change) < -0.25) are defined by comparing patient and control within each cell type. **(C)** Boxplots show bulk RNA-seq signal as $log_2$(fold change) of fragments per kilobase per million reads ($log_2$(FPKM)) of differentially expressed genes defined by snRNA-seq for all samples. Each dot represents one sample. **(D)** Heatmaps showing bulk ATAC-seq signal of increased (top left) and decreased peaks (bottom left) between patient and control samples compared to shuffle (right) controls. The color scale indicates reads per kilobase per million reads (RPKM) calculated z-score. **(E)** Volcano plots showing differential chromatin accessibility of all 8 cell types as defined from snATAC-seq. Increased peaks (red) (FDR <0.1 and $log_2$ (fold change) > 1) and decreased peaks (blue) (FDR <0.1 and $log_2$ (fold change) < -1) are defined by comparing patient and control within each cell type. **(F)** HOMER (*96*) motif analysis of differentially accessible peaks reveals significant enrichment of sequences at both increased (right) and decreased (left) peaks, as measured by the bulk ATAC-seq. **(G)** A genome browser screenshot shows the upregulation of the *DNMT1* gene with pseudo-bulk tracks of Syncytiotrophoblast (ST) cells from snRNA-seq. Signals are displayed as Reads Per Million mapped reads (RPM) and the y-axes are between 0 and 2 for all tracks.

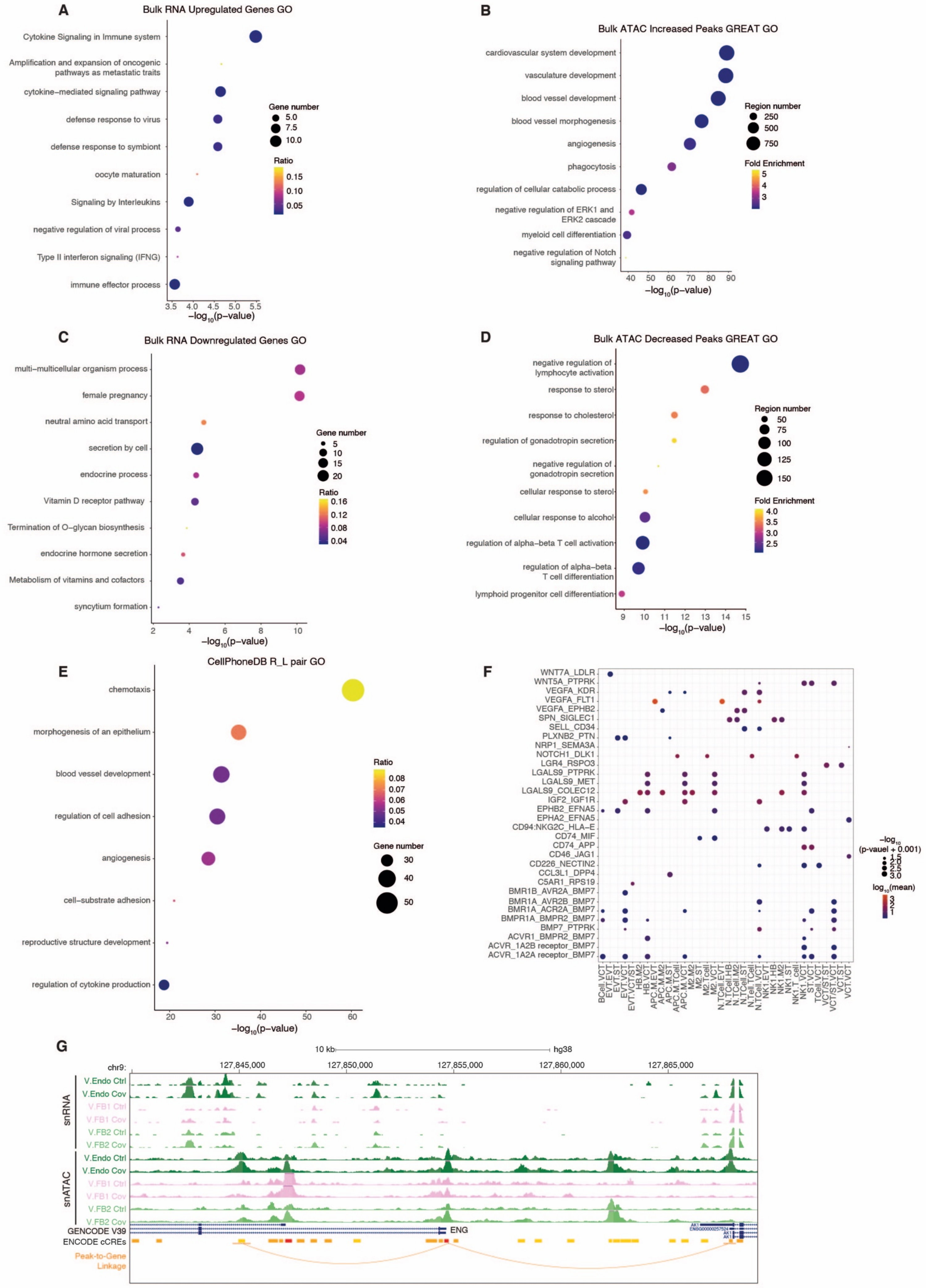

A
Bulk RNA Upregulated Genes GO
Cytokine Signaling in Immune system
Amplification and expansion of oncogenic pathways as metastatic traits
cytokine-mediated signaling pathway
defense response to virus
defense response to symbiont
oocyte maturation
Signaling by Interleukins
negative regulation of viral process
Type II interferon signaling (IFNG)
immune effector process
−log10(p−value)
Gene number
Ratio
B
Bulk ATAC Increased Peaks GREAT GO
cardiovascular system development
vasculature development
blood vessel development
blood vessel morphogenesis
angiogenesis
phagocytosis
regulation of cellular catabolic process
negative regulation of ERK1 and ERK2 cascade
myeloid cell differentiation
negative regulation of Notch signaling pathway
Region number
Fold Enrichment
C
Bulk RNA Downregulated Genes GO
multi−multicellular organism process
female pregnancy
neutral amino acid transport
secretion by cell
endocrine process
Vitamin D receptor pathway
Termination of O−glycan biosynthesis
endocrine hormone secretion
Metabolism of vitamins and cofactors
syncytium formation
D
Bulk ATAC Decreased Peaks GREAT GO
negative regulation of lymphocyte activation
response to sterol
response to cholesterol
regulation of gonadotropin secretion
negative regulation of gonadotropin secretion
cellular response to sterol
cellular response to alcohol
regulation of alpha-beta T cell activation
regulation of alpha−beta T cell differentiation
lymphoid progenitor cell differentiation
E
CellPhoneDB R_L pair GO
chemotaxis
morphogenesis of an epithelium
blood vessel development
regulation of cell adhesion
angiogenesis
cell−substrate adhesion
reproductive structure development
regulation of cytokine production
F
−log10 (p-vauel + 0.001)
log10(mean)
G
snRNA
snATAC
V.Endo Ctrl
V.Endo Cov
V.FB1 Ctrl
V.FB1 Cov
V.FB2 Ctrl
V.FB2 Cov
GENCODE V39
ENCODE cCREs
Peak-to-Gene Linkage
ENG
hg38
chr9:

**Fig. S4. Gene Ontology (GO) analysis and CellphoneDB results for bulk assays and snRNA-seq. (A-D)** Bubble plots showing GO analysis with Metascape (*97*) for **(A)** upregulated genes in bulk RNA-seq, **(B)** increased peaks in bulk ATAC-seq, **(C)** downregulated genes in bulk RNA-seq, and **(D)** decreased peaks in bulk ATAC-seq. For the bubble plots derived from the bulk RNA-seq, the size and color of the bubbles represent the number of dysregulated genes and the ratio to total genes under each GO term, respectively. For the bubble plots derived from the bulk ATAC-seq, the size and color of the bubbles represent the number of differential regions and the fold enrichment of peaks in each category. **(E)** Bubble plot showing GO analysis of significant, patient specific receptor-ligands interactions defined by CellPhoneDB (*42*), in which the ligand is differentially expressed in given cell types. The size and color of the bubble represent the number of dysregulated genes and the ratio to total genes under each GO term, respectively. **(F)** Bubble plot of significant, patient specific receptor-ligand pairs under the GO term of "regulation of cytokine production". The interactions between immune and placenta cell types are shown. The color of the bubbles indicates the $log_{10}$ transformed mean, and the size of the bubbles shows the significance through negative $log_{10}$(p-value). **(G)** A genome browser screenshot of the region containing the *ENG* gene. The snRNA-seq and snATAC-seq pseudo-bulk tracks of Vilious Endothelial cells (V.Endo), Vilious Fibroblasts 1 (V.FB1), and Villous Fibroblasts 2 (V.FB2) of patients (Cov) and controls (Ctrl) are displayed as RPM. The screenshot also includes the ENCODE cCREs and the ArchR (*34*) peak-to-gene linkage tracks (orange arc). The y-axes are between 0 and 2 for all tracks.

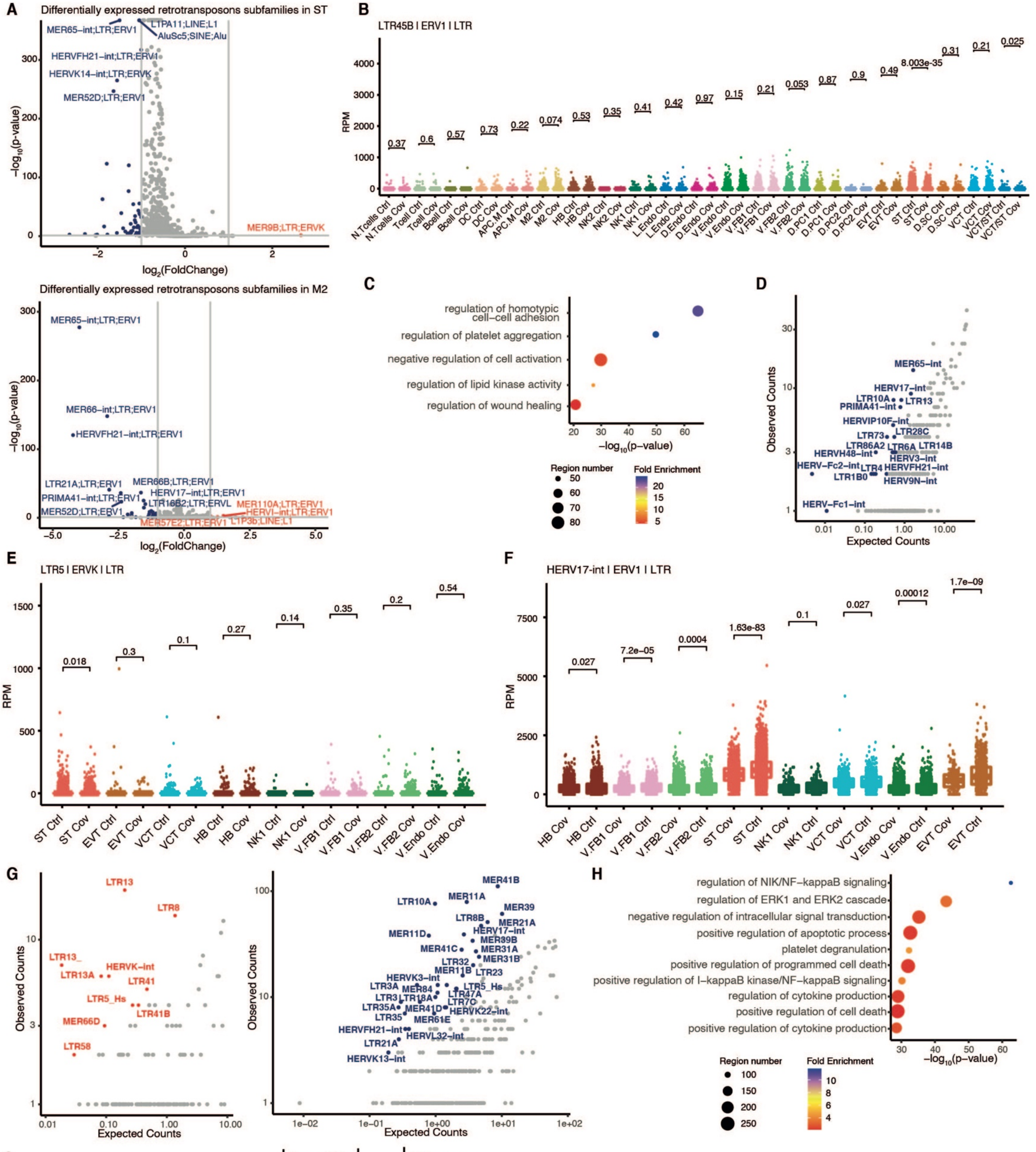


I

| Motif | Transcription factor | P-value | # Sequence with motif |
|---|---|---|---|
| 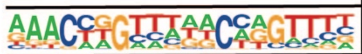 | GRHL2 | 1e-5 | 65 |
| 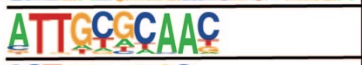 | CEBP | 1e-4 | 129 |
| 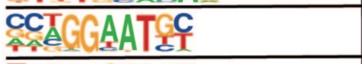 | TEAD4 | 1e-4 | 117 |
| 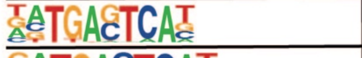 | BATF | 1e-4 | 89 |
| 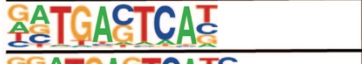 | JunB | 1e-3 | 73 |
| 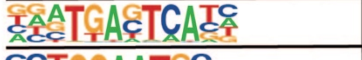 | Fra1 | 1e-3 | 72 |
| 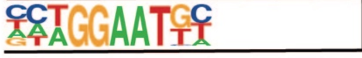 | TEAD | 1e-2 | 97 |

**Fig S5. Dysregulation of retrotransposons in SARS-CoV-2 infected pregnancies. (A)** Volcano plot showing differentially expressed retrotransposons subfamilies between control and patient in ST (top) and M2 Macrophage (M2) cells (bottom) as measured by snRNA-seq. For each subfamily, the negative $\log_{10}$ (padj) is plotted against the $\log_2$ (fold change) of RPKM (patient/control). Significantly upregulated (p-value < 0.1 and $\log_2$ (fold change) > 1 ; n = 1 (ST) and 4 (M2)) and downregulated subfamilies (p-value < 0.1 and $\log_2$ (fold change) < -1 ; n = 48 (ST) and 30 (M2)) are labelled with red and blue, respectively. **(B)** Boxplot showing the expression of LTR45B subfamily in distinct cell types as determined by snRNA-seq, showing significant downregulation only in ST cells. Each dot represents one nucleus and the p-value represents significance calculated by student T-test. **(C)** Genomic Regions Enrichment of Annotations Tool (GREAT) (*98*) analysis of significantly upregulated individual retrotransposons in patient samples as measured by bulk RNA-seq. **(D)** Comparison of observed versus expected counts of significantly downregulated individual retrotransposons within subfamilies as determined by bulk RNA-seq. Subfamilies with observed/expected count > 5 and p-value < 0.001 are labelled in blue. **(E-F)** Boxplots show the chromatin accessibility of **(E)** LTR5 subfamily and **(F)** HERV17-int subfamily in distinct cell types as determined by snATAC-seq. Each dot represents one nucleus and the p-value represents significance as calculated by student T-test. **(G)** Comparison of observed versus expected numbers of individual retrotransposons with significantly increased (left) and decreased (right) chromatin accessibility within subfamilies as determined by bulk ATAC-seq. Subfamilies with observed/expected count > 5 and p-value < 0.001 are labelled with red (left) and blue (right). **(H)** GREAT analysis of individual retrotransposons with increased chromatin accessibility from bulk ATAC-seq shows enrichment of GO terms including immune regulation. **(I)** HOMER motif analysis of individual retrotransposons with decreased chromatin accessibility in ST cells as determined by snATAC-seq.

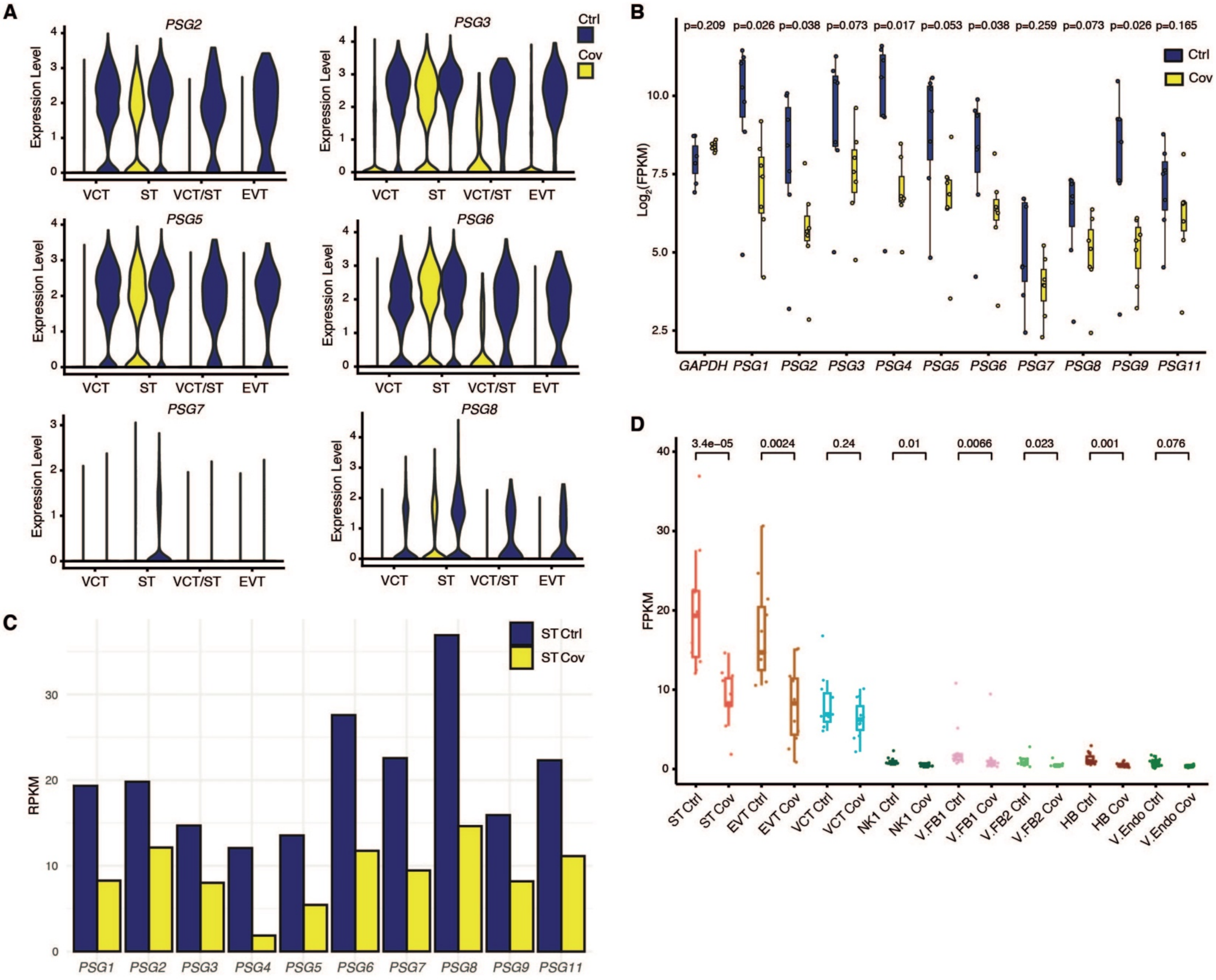

A
PSG2
PSG3
PSG5
PSG6
PSG7
PSG8
Ctrl
Cov
Expression Level
VCT
ST
VCT/ST
EVT
B
p=0.209 p=0.026 p=0.038 p=0.073 p=0.017 p=0.053 p=0.038 p=0.259 p=0.073 p=0.026 p=0.165
Log2(FPKM)
GAPDH PSG1 PSG2 PSG3 PSG4 PSG5 PSG6 PSG7 PSG8 PSG9 PSG11
C
ST Ctrl
ST Cov
RPKM
PSG1 PSG2 PSG3 PSG4 PSG5 PSG6 PSG7 PSG8 PSG9 PSG11
D
3.4e−05 0.0024 0.24 0.01 0.0066 0.023 0.001 0.076
FPKM
ST Ctrl ST Cov EVT Ctrl EVT Cov VCT Ctrl VCT Cov NK1 Ctrl NK1 Cov V.FB1 Ctrl V.FB1 Cov V.FB2 Ctrl V.FB2 Cov HB Ctrl HB Cov V.Endo Ctrl V.Endo Cov

**Fig. S6. Downregulation of *PSG* genes' expression in SARS-CoV-2 infected pregnancies. (A)** Violin plots show decreased expression of *PSG2* gene (top left), *PSG3* gene (top right), *PSG5* gene (middle left), *PSG6* gene (middle right), *PSG7* gene (bottom left), and *PSG8* gene (bottom right) in respective trophoblast cells from snRNA-seq in patient and control samples. Expression values are displayed as normalized counts. **(B)** Box plot showing decreased expression of *PSG* genes in patient versus controls as measured by bulk RNA-seq. Each dot represents a sample and the p-value represents the significance as calculated by student T-test. **(C)** Bar chart showing aggregated RPKM signal of each intronic LTR8B at a *PSG* gene in ST cells, as determined by snATAC-seq. **(D)** Boxplot showing decreased aggregated chromatin accessibility signal of *PSG* genes intronic LTR8B elements in distinct cell types as determined by snATAC-seq. Each dot represents one LTR8B element and the p-value represents significance as calculated by the Wilcoxon test.

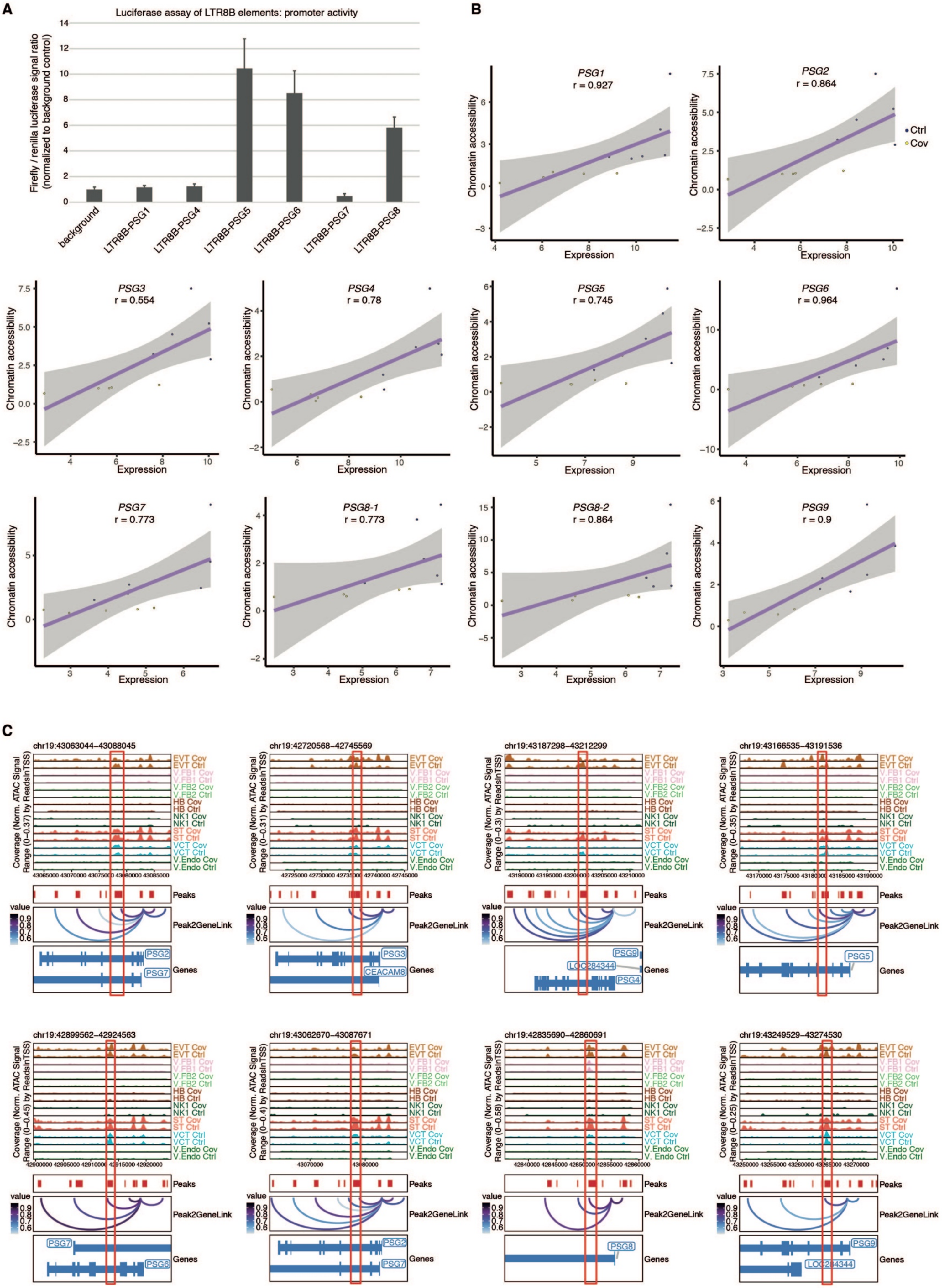

A
Luciferase assay of LTR8B elements: promoter activity
Firefly / renilla luciferase signal ratio (normalized to background control)
background
LTR8B-PSG1
LTR8B-PSG4
LTR8B-PSG5
LTR8B-PSG6
LTR8B-PSG7
LTR8B-PSG8
B
PSG1
r = 0.927
PSG2
r = 0.864
PSG3
r = 0.554
PSG4
r = 0.78
PSG5
r = 0.745
PSG6
r = 0.964
PSG7
r = 0.773
PSG8-1
r = 0.773
PSG8-2
r = 0.864
PSG9
r = 0.9
Chromatin accessibility
Expression
Ctrl
Cov
C
chr19:43063044-43088045
chr19:42720568-42745569
chr19:43187298-43212299
chr19:43166535-43191536
chr19:42899562-42924563
chr19:43062670-43087671
chr19:42835690-42860691
chr19:43249529-43274530
Coverage (Norm. ATAC Signal Range (0–0.37) by ReadsInTSS)
EVT Cov
EVT Ctrl
V.FB1 Cov
V.FB1 Ctrl
V.FB2 Cov
V.FB2 Ctrl
HB Cov
HB Ctrl
NK1 Cov
NK1 Ctrl
ST Cov
ST Ctrl
VCT Cov
VCT Ctrl
V.Endo Cov
V.Endo Ctrl
Peaks
Peak2GeneLink
value
Genes
PSG2
PSG7
PSG3
CEACAM8
PSG9
LOC284344
PSG4
PSG5
PSG6
PSG8

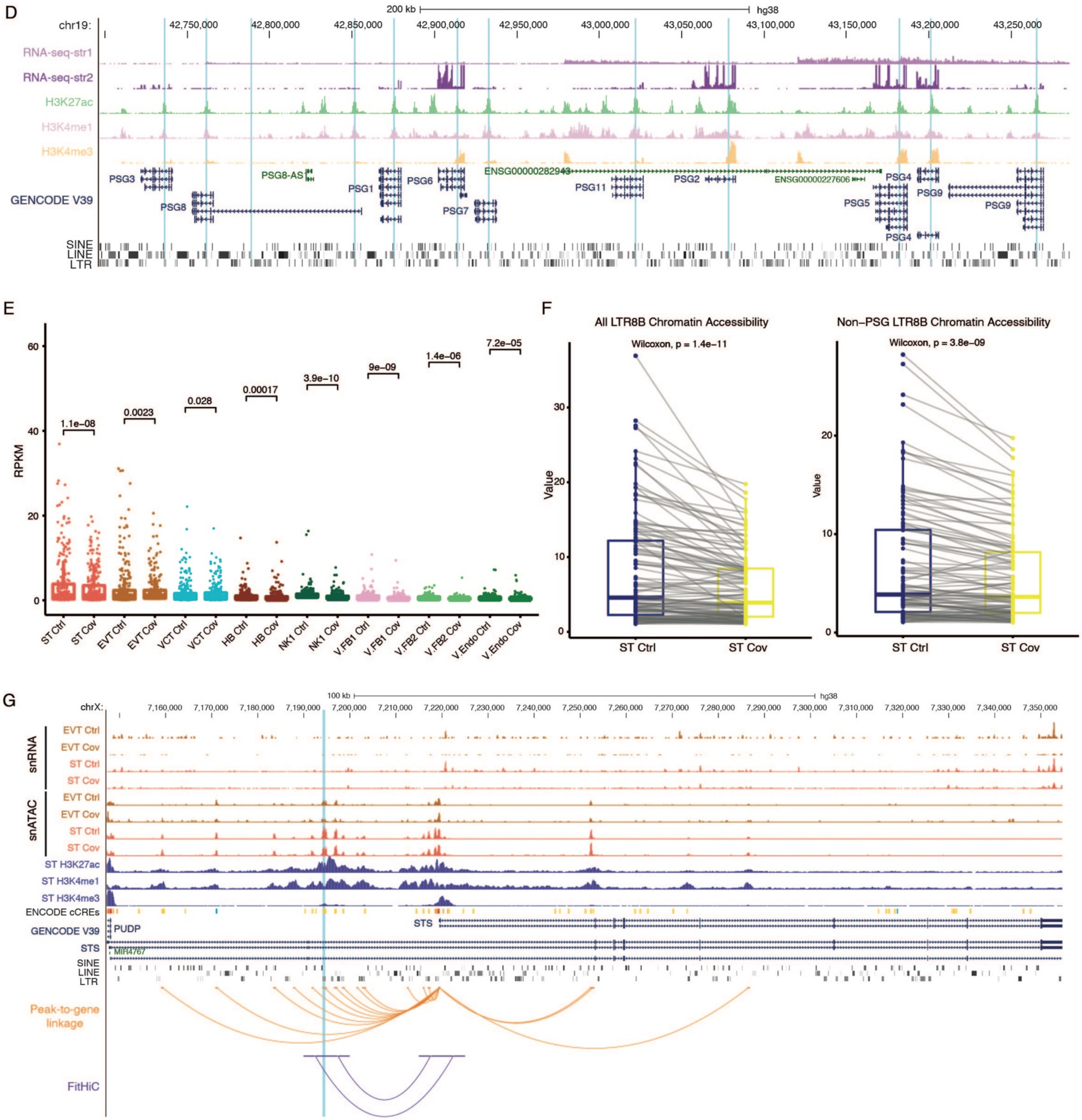
D
200 kb
hg38
chr19:
42,750,000
42,800,000
42,850,000
42,900,000
42,950,000
43,000,000
43,050,000
43,100,000
43,150,000
43,200,000
43,250,000
RNA-seq-str1
RNA-seq-str2
H3K27ac
H3K4me1
H3K4me3
GENCODE V39
PSG3
PSG8
PSG8-AS
PSG1
PSG6
PSG7
ENSG00000282943
PSG11
PSG2
ENSG00000227606
PSG4
PSG5
PSG9
SINE
LINE
LTR
E
RPKM
60
40
20
0
1.1e−08
0.0023
0.028
0.00017
3.9e−10
9e−09
1.4e−06
7.2e−05
ST Ctrl
ST Cov
EVT Ctrl
EVT Cov
VCT Ctrl
VCT Cov
HB Ctrl
HB Cov
NK1 Ctrl
NK1 Cov
V.FB1 Ctrl
V.FB1 Cov
V.FB2 Ctrl
V.FB2 Cov
V.Endo Ctrl
V.Endo Cov
F
All LTR8B Chromatin Accessibility
Wilcoxon, p = 1.4e−11
Non−PSG LTR8B Chromatin Accessibility
Wilcoxon, p = 3.8e−09
Value
30
20
10
0
ST Ctrl
ST Cov
G
100 kb
hg38
chrX:
7,160,000
7,170,000
7,180,000
7,190,000
7,200,000
7,210,000
7,220,000
7,230,000
7,240,000
7,250,000
7,260,000
7,270,000
7,280,000
7,290,000
7,300,000
7,310,000
7,320,000
7,330,000
7,340,000
7,350,000
snRNA
snATAC
EVT Ctrl
EVT Cov
ST Ctrl
ST Cov
ST H3K27ac
ST H3K4me1
ST H3K4me3
ENCODE cCREs
GENCODE V39
PUDP
STS
MIR4767
SINE
LINE
LTR
Peak-to-gene linkage
FitHiC

**Fig. S7. Decreased expression of *PSG* genes is associated with reduced intronic LTR8B chromatin accessibility. (A)** Luciferase assay of individual LTR8B elements within each *PSG* gene shows the sufficiency of particular elements to serve as promoters. Y-axis value represents Firefly/Renilla luciferase signal ratio normalized to background control. **(B)** Dot plots showing correlations of the transcriptional levels of *PSG*s and the ATAC-seq signals of corresponding intronic LTR8B elements as measured by bulk RNA-seq and ATAC-seq, respectively. X-axis is displayed as $\log_2$(FPKM) as measured by bulk RNA-seq and the y-axis is displayed as RPKM as determined by bulk ATAC-seq. Spearman correlation is used to calculate the correlation. **(C)** Peak-to-gene linkage analysis by ArchR of *PSG* genes and their corresponding intronic LTR8B elements (red box) suggests the potential interactions between gene promoters and their corresponding intronic LTR8B element. Datasets are displayed as aggregated normalized ATAC-seq signals of distinct cell types as measured by snATAC-seq. **(D)** A genome browser screenshot showing expression of *PSG* genes and enrichment patterns of active histone modifications (ChIP-seq for H3K27ac, H3K4me1, and H3K4me3) in Trophoblast Stem Cell (TSC). All datasets are displayed as RPM, with y-axis ranging from 0 to 2 for RNA-seq, 0 to 80 for H3K27ac ChIP-seq, 0 to 40 for H3K4me1 ChIP-seq, and 0 to 80 for H3K4me3. The blue shading highlights the intronic LTR8B elements within *PSG* genes. **(E)** Boxplot shows the aggregated ATAC-seq signal (RPKM) of all elements of the LTR8B subfamily in distinct cell types as determined by snATAC-seq. P-value represents significance as calculated by Wilcoxon test. Each dot represents one individual LTR8B element. **(F)** Parallel coordinate plots displaying the significantly decreased chromatin accessibility of (left) all LTR8B elements (n=118) and (right) LTR8B elements outside the *PSG* cluster (n=107) upon SARS-CoV-2 infection. To reduce the contribution of background signal, LTR8B elements are filtered by RPKM > 1 in either control or patient sample and length > 400bp. Significance is calculated by Wilcoxon test. **(G)** A genome browser screenshot shows an LTR8B element with subtle decreased chromatin accessibility (blue shading) in trophoblast cells interacting with *STS* promoter. Concomitantly the expression of *STS* is downregulated. Orange arcs represent interactions defined from Peak-to-gene linkage analysis and purple arcs represent interactions defined from Fit-Hi-C analysis. All datasets are displayed as RPM, with y-axis ranging from 0 to 20 for snRNA-seq, 0 to 5 for snATAC-seq, 0 to 200 for H3K27ac ChIP-seq, 0 to 200 for H3K4me1 ChIP-seq, and 0 to 500 for H3K4me3.

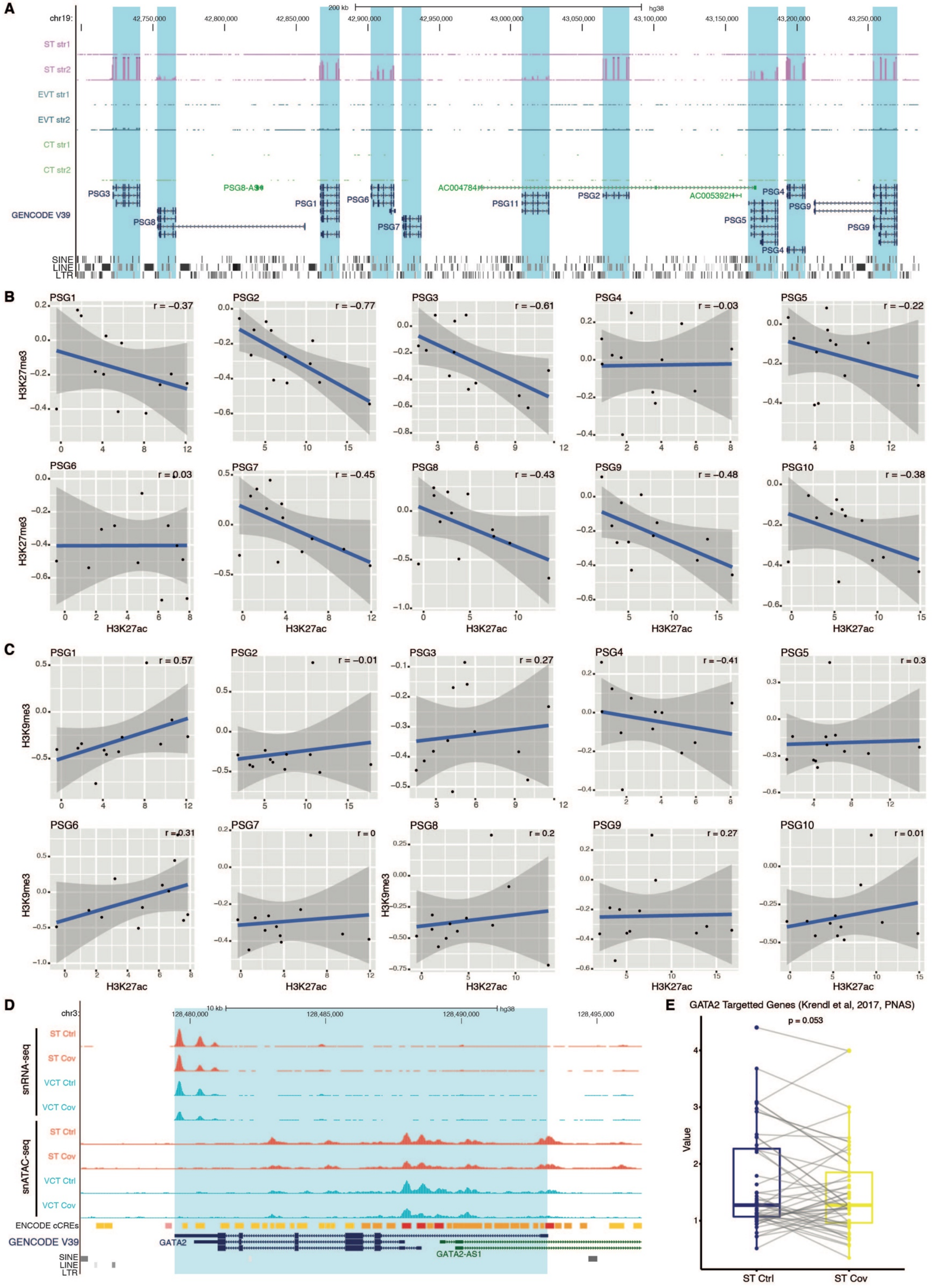
A
chr19:
hg38
200 kb
ST str1
ST str2
EVT str1
EVT str2
CT str1
CT str2
GENCODE V39
SINE
LINE
LTR
B
H3K27me3
H3K27ac
C
H3K9me3
H3K27ac
D
chr3:
10 kb
snRNA-seq
snATAC-seq
ST Ctrl
ST Cov
VCT Ctrl
VCT Cov
ENCODE cCREs
GENCODE V39
GATA2
GATA2-AS1
E
GATA2 Targetted Genes (Krendl et al, 2017, PNAS)
p = 0.053
Value
ST Ctrl
ST Cov

**Fig. S8. Epigenetic regulation of PSG genes in trophoblasts and the downregulation of GATA2 and its targets in patient samples. (A)** A genome browser screenshot shows the expression of the *PSG* gene cluster in ST, Extravilious Trophoblast (EVT), and Cytotrophoblast (CT) cell types. RNA-seq datasets from primary trophoblasts are obtained from the International human epigenome consortium (IHEC) public data repository. RNA-seq tracks are displayed as RPM, with y-axis ranging from 0 to 1000. **(B-C)** Dot plots show the correlation of **(B)** H3K27me3 versus H3K27ac and **(C)** H3K9me3 versus H3K27ac at *PSG* intronic LTR8B in ST, EVT and CT cells. ChIP-seq datasets from primary trophoblasts are obtained from the IHEC public data repository. Each plot represents one PSG intronic LTR8B and each dot represents a biological sample. Spearman correlation coefficient is calculated. **(D)** A genome browser screenshot shows decreased expression and chromatin accessibility of the *GATA2* gene in ST and Vilious Cytotrophoblast (VCT) cells as measured by snRNA-seq and snATAC-seq. Aggregated snRNA-seq and snATAC-seq datasets are displayed as RPM, with y-axis ranging from 0 to 40 and 0 to 5, respectively. **(E)** Boxplot shows the expression of *GATA2* targeted genes (n=49) (*63*) in control and patient samples in ST cells as measured by snRNA-seq. Values on the y-axis are displayed as RPKM values.

**Table S1. Summary of the samples used in this study**

| Characteristics | Cases<br>n=7 | Controls<br>n=7 |
|---|---|---|
| Maternal age (years) | 35 (27-42) | 37 (33-40) |
| Ethnicity | | |
| Chinese | 4 (57.1%) | 7 (100%) |
| Non-Chinese Asian | 3 (42.9%) | 0 (0%) |
| Nulliparous | 1 (14.3%) | 0 (0%) |
| Prepregnancy body mass index ($kg/m^2$) | 22.8 (18.0-35.3) | 21.1 (18.8-25.9) |
| Diabetes (preexisting or gestational) | 0 (0%) | 1 (14.3%) |
| Cesarean delivery | 4 (57.1%) | 7 (100%) |
| GA at delivery in weeks | 37.0 (32.4-40.4) | 38.1 (37-38.6) |
| Preterm delivery at GA <37 weeks | 3 (42.9%) | 0 (0%) |
| Infant sex: male | 2 (28.6%) | 1 (14.3%) |
| Birthweight (grams) | 3075 (1890-3760) | 2960 (2525-3320) |
| Symptomatic | 5 (71.4%) | - |
| GA at symptom onset (weeks) | 35.6 (31.3-39.0) | - |
| GA at diagnosis (weeks) | 35.9 (31.6-39.6) | - |
| GA at hospital admission (weeks) | 35.7 (31.6-39.6) | - |
| Ct value at diagnosis | 25.4 (12.5-34.2) | - |
| Antigen positive duration (weeks) | 1.9 (0.7-4) | - |
| Infection-to-delivery interval (weeks) | 0.6 (0.1-8.1) | - |
| Seroconversion interval (weeks) | 1.7 (0.6-2.4) | - |
| Maternal IgG seropositive at delivery | 3 (42.9%) | - |
| Maternal IgG concentrations at delivery (ng/ml) | 306.5 (291.6-697.8) | - |
| Maternal IgM seropositive at delivery | 3 (42.9%) | - |
| Maternal IgM concentrations at delivery (ng/ml) | 47.3 (5.6-57.0) | - |
| Infant IgG seropositive at delivery | 1 (14.3%) | - |
| Infant IgG concentrations at delivery (ng/ml) | 777.4 | - |
| Transfer ratio at delivery | 1.11* | - |

Numerical variables presented in median (range)
Categorical variables presented in number (%)
GA=gestational age; Ct=cycle threshold values in qPCR
*3 mothers had anti-SARS-CoV-2 IgG at delivery but only 1 infant had IgG at birth.

**Table S2. Summary of the COVID-19 status of patients**

| Case | Date of symptom onset | GA of symptom onset | Test positive date | Test positive GA | EDD | Date of hospital admission | Date of delivery | Date of hospital discharge | Test negative date | Test negative GA |
|---|---|---|---|---|---|---|---|---|---|---|
| Cov1 | 18-Jul-20 | 34W0D | 21-Jul-20 | 34W3D | 29-Aug-20 | 21-Jul-20 | 22-Jul-20 | 18-Aug-20 | 17/8/2020 | PP day 26 |
| Cov2 | 25-Aug-20 | 39W0D | 29-Aug-20 | 39W4D | 1-Sep-20 | 29-Aug-20 | 2-Sep-20 | 16-Sep-20 | No negative result | No negative result |
| Cov3 | Asymptomatic | Asymptomatic | 17-Aug-20 | 32W2D | 10-Oct-20 | 18-Aug-20 | 13-Oct-20 | 4-Sep-20 | 3-Sep-20 | 34W5D |
| Cov4 | Asymptomatic | Asymptomatic | 9-Dec-20 | 39W4D | 12-Dec-20 | 9-Dec-20 | 10-Dec-20 | 23-Dec-20 | No negative result | No negative result |
| Cov5 | 16-Dec-20 | 35W4D | 18-Dec-20 | 35W6D | 16-Jan-21 | 17-Dec-20 | 22-Dec-20 | 5-Jan-21 | No negative result | No negative result |
| Cov6 | 19-Dec-20 | 31W2D | 21-Dec-20 | 31W4D | 18-Feb-21 | 21-Dec-20 | 27-Dec-20 | 6-Jan-21 | No negative result | No negative result |
| Cov7 | 24-Jan-21 | 36W3D | 24-Jan-21 | 36W3D | 18-Feb-21 | 24-Jan-21 | 28-Jan-21 | 8-Feb-21 | No negative result | No negative result |

GA=gestational age

**Table S3. RT-qPCR for the SARS-CoV-2 Nucleocapsid protein for each sample.**

| | Mean CT value of duplicate | | |
|---|---|---|---|
| | N1 | N2 | RP |
| Positive Control | 28.82 | 30.12 | 30.18 |
| NTC | Undetermined | Undetermined | Undetermined |
| 1000 copies/ reaction | 24.82 | 25.61 | Undetermined |
| 100 copies/ reaction | 28.54 | 29.36 | Undetermined |
| 10 copies/ reaction | 31.90 | 32.77 | Undetermined |
| Cov1-1 | Undetermined | Undetermined | 18.10 |
| Cov1-2 | Undetermined | Undetermined | 18.36 |
| Cov1-3 | Undetermined | Undetermined | 17.89 |
| Cov2-1 | Undetermined | Undetermined | 18.26 |
| Cov2-2 | Undetermined | Undetermined | 18.10 |
| Cov2-3 | Undetermined | Undetermined | 18.32 |
| Cov3-1 | Undetermined | Undetermined | 17.98 |
| Cov3-2 | Undetermined | Undetermined | 17.78 |
| Cov3-3 | Undetermined | Undetermined | 17.84 |
| Cov4-1 | Undetermined | Undetermined | 18.09 |
| Cov4-2 | Undetermined | Undetermined | 18.26 |
| Cov4-3 | Undetermined | Undetermined | 18.19 |
| Cov5-1 | Undetermined | Undetermined | 18.54 |
| Cov5-2 | Undetermined | Undetermined | 18.51 |
| Cov5-3 | Undetermined | Undetermined | 18.35 |
| Cov6-1 | Undetermined | Undetermined | 18.96 |
| Cov6-2 | Undetermined | Undetermined | 18.33 |
| Cov6-3 | Undetermined | Undetermined | 18.15 |
| Cov7-1 | Undetermined | Undetermined | 17.79 |
| Cov7-2 | Undetermined | Undetermined | 17.92 |
| Cov7-3 | Undetermined | Undetermined | 18.20 |

**Table S4. Summary of assays conducted on control and patient samples in this study.**

| Sample | Bulk RNA-seq | Bulk ATAC-seq | snRNA-seq | snATAC-seq |
|---|---|---|---|---|
| **Ctrl1** | Yes | Yes | / | Yes |
| **Ctrl2** | Yes | Yes | Yes | Yes |
| **Ctrl3** | Yes | Yes | / | / |
| **Ctrl4** | Yes | / | / | / |
| **Ctrl5** | Yes | Yes | Yes | / |
| **Ctrl6** | Yes | Yes | / | / |
| **Ctrl7** | Yes | Yes | Yes | / |
| **Cov1** | Yes | / | / | / |
| **Cov2** | Yes | / | Yes | Yes |
| **Cov3** | Yes | Yes | / | Yes |
| **Cov4** | Yes | Yes | Yes | Yes |
| **Cov5** | Yes | Yes | Yes | / |
| **Cov6** | Yes | Yes | / | / |
| **Cov7** | Yes | Yes | / | / |

### Table S5. List of primers used for this study.

| Primer name | Primer type | Assay | Primer sequence |
|---|---|---|---|
| PSG1-RT-F | qPCR primer | RT-qPCR for checking PSG expression | GAGGAGTAACTGGACGTTTCACC |
| PSG1-RT-R | qPCR primer | RT-qPCR for checking PSG expression | TGGAGTCTCAGGGTCACAGGTT |
| PSG2-RT-F | qPCR primer | RT-qPCR for checking PSG expression | AGTGACCCAGTCACCCTGAATC |
| PSG2-RT-R | qPCR primer | RT-qPCR for checking PSG expression | CGAAGCAAGACAAGTAGAGG |
| PSG3-RT-F | qPCR primer | RT-qPCR for checking PSG expression | CGTAAAGCGAGGTGATGGGACT |
| PSG3-RT-R | qPCR primer | RT-qPCR for checking PSG expression | AAGCTCACAGCCTCCATGTCCT |
| PSG4-RT-F | qPCR primer | RT-qPCR for checking PSG expression | CCCAGTCACCCTGAATG |
| PSG4-RT-R | qPCR primer | RT-qPCR for checking PSG expression | ATTGTGCCCGTGGGTTAGACTC |
| PSG5-RT-F | qPCR primer | RT-qPCR for checking PSG expression | CGGTGGGTTAGATTCCGC |
| PSG5-RT-R | qPCR primer | RT-qPCR for checking PSG expression | TCACCCTGAATGTCCTCTGT |
| PSG7-RT-F | qPCR primer | RT-qPCR for checking PSG expression | AAAGCGAGGTGATGGGACTGGA |
| PSG7-RT-R | qPCR primer | RT-qPCR for checking PSG expression | CTGGAGTCTCAGGATCACAGGT |
| PSG8-RT-F | qPCR primer | RT-qPCR for checking PSG expression | TCCAAGAATACTGTGCCG |
| PSG8-RT-R | qPCR primer | RT-qPCR for checking PSG expression | ATCCGCAGTTACCCAGTCAC |
| PSG9-RT-F | qPCR primer | RT-qPCR for checking PSG expression | TAGGTAGCTTGCGTCCAGAG |
| PSG9-RT-R | qPCR primer | RT-qPCR for checking PSG expression | GATCCAGAATGTCACCCG |
| LTR8B-PSG8-1-F | PCR primer | Amplify LTR8B sequence for luciferase assay | cgcggatccGGTGAGTCAGAGCAAAGATTACA |
| LTR8B-PSG8-1-R | PCR primer | Amplify LTR8B sequence for luciferase assay | cgcggatccACAGGGATAGGCAGTTCTCCA |
| LTR8B-PSG1-F | PCR primer | Amplify LTR8B sequence for luciferase assay | cgcggatccCTGGTGAGTCAGTGCAGAGA |
| LTR8B-PSG1-R | PCR primer | Amplify LTR8B sequence for luciferase assay | cgcggatccGGCAGTTCCTCCATCCACTC |
| LTR8B-PSG6-F | PCR primer | Amplify LTR8B sequence for luciferase assay | cgcggatccGGGTGAAATGAGCCTATGGG |
| LTR8B-PSG6-R | PCR primer | Amplify LTR8B sequence for luciferase assay | cgcggatccCAAGTTCCTCTGGCCTCCTG |
| LTR8B-PSG7-F | PCR primer | Amplify LTR8B sequence for luciferase assay | cgcggatccATGGGCTTTGGGAACTGC |
| LTR8B-PSG7-R | PCR primer | Amplify LTR8B sequence for luciferase assay | cgcggatccTCCTGCACACATACATACACCA |
| LTR8B-PSG5-F | PCR primer | Amplify LTR8B sequence for luciferase assay | cgcggatccACAACAGTGACAGCAAAGTAGC |
| LTR8B-PSG5-R | PCR primer | Amplify LTR8B sequence for luciferase assay | cgcggatccGGCAGTTCCTCCATCCATTCA |
| LTR8B-PSG4-F | PCR primer | Amplify LTR8B sequence for luciferase assay | cgcggatccCTGGTGAGTCAGTGCAAAGAT |
| LTR8B-PSG4-R | PCR primer | Amplify LTR8B sequence for luciferase assay | cgcggatccGGCAGTTCCTCCATCCACTC |
| GFP_luciferase_F | PCR primer | Amplify GFP sequence for luciferase assay | ggatccCACATGAAGCAGCACGACTT |
| GFP_luciferase_R | PCR primer | Amplify GFP sequence for luciferase assay | ggatccTCCATGCCGAGAGTGATCC |
